# The microbiome determines the phenotype in CTLA-4 insufficient mice and men

**DOI:** 10.64898/2026.08.14.742114

**Authors:** Bei Zhao, Lara Stopp, Pavaret Sivapornnukul, Simon Lagies, Kun D. Huang, Till Robin Lesker, Virginia Andreani, Rabina Giri, Pavla Mrovecova, Wiebke Schifferdecker, Angelika Hofmann, Katja Gräwe, Luis Braun, Jakob Begun, Christoph Schell, Stephan P. Rosshart, Bernd Kammerer, Till Strowig, Bodo Grimbacher

## Abstract

CTLA-4 (haplo)insufficiency displays incomplete penetrance and phenotypic heterogeneity, indicating the involvement of additional disease modifiers beyond the genetic defect. Microbiome analyses reveal a positive association between disease severity and intestinal dysbiosis, highlighting the microbiome as a critical contributor. To investigate this relationship mechanistically, we generated *Ctla4⁺^/^⁻* wildlings harboring a natural microbiota. Unlike specific pathogen free (SPF) counterparts, which remain healthy, *Ctla4⁺^/^⁻* wildlings spontaneously develop disease phenotypes resembling human CTLA-4 haploinsufficiency. Disease onset is followed by reduced microbial diversity and expansion of pathobionts. Integrative immunophenotyping shows that the natural microbiota synergizes with *Ctla4* haploinsufficiency to reshape innate and adaptive immune compartments, generating a sustained pro-inflammatory milieu and reduced CTLA-4 expression in the cecum. Furthermore, microbiota-derived metabolites promote inflammatory cytokine production in both murine and human primary T cells via NF-κB activation. Collectively, *Ctla4⁺^/^⁻* wildlings constitute an effective model for dissecting microbiome–immune crosstalk in CTLA-4 (haplo)insufficiency and for exploring therapeutic strategies.

## Main Text

Cytotoxic T-lymphocyte antigen-4 (CTLA-4) is a key inhibitory receptor restraining T-cell activation and preserving peripheral immune tolerance. Its immunoregulatory function, first identified in 1987 (*1*), has since been extensively characterized, laying the foundation for successful clinical translation in cancer immunotherapy. Blockade of CTLA-4 with agents such as ipilimumab enhances CD28-mediated co-stimulation and promotes antitumor immunity (*2*). However, therapeutic CTLA-4 inhibition can also disrupt immune homeostasis, leading to immune-related adverse events (irAEs) – most notably enterocolitis – which occur in approximately 60-70% of treated individuals (*3*). This observation highlights the essential role of CTLA-4 in maintaining mucosal tolerance.

A naturally occurring human monogenic condition mirrors these therapeutic toxicities: CTLA-4 *haplo*insufficiency may be caused by monoallelic nonsense mutations in *CTLA4*, but missense mutations in *CTLA4* may impair ligand binding or dimerization of CTLA-4, thereby impairing CTLA-4 biology, leading to so called “CTLA-4 insufficiency”. Since its identification as a monogenic immune dysregulation disorder in 2014 (*4, 5*), CTLA-4 (haplo)insufficiency has been recognized as a multisystem disease characterized by hypogammaglobulinemia, recurrent infections, autoimmune cytopenias, lymphoproliferation, and enteropathy closely resembling irAEs observed after therapeutic CTLA-4 blockade (*6*). Although inherited in an autosomal-dominant manner, the condition exhibits striking phenotypic heterogeneity: some variant carriers remain asymptomatic, whereas others develop severe, even life-threatening immune dysregulation. With an estimated penetrance of ∼70% (*7*), additional non-genetic modifiers are likely to contribute to disease penetrance and expressivity. Hence, identifying these modifiers is essential not only for accurate diagnosis, risk prediction, and targeted therapeutic intervention in patients with CTLA-4 (haplo)insufficiency but also for patients with irAEs during checkpoint blockade. Among potential environmental modifiers, the gut microbiome has emerged as a particularly compelling candidate. Recent work has shown that alterations in the microbiome are associated with disease severity in several monogenic immune disorders (*8, 9*), including CTLA-4 (haplo)insufficiency (*10*). Moreover, the microbial content also appears to affect side effects during checkpoint blockade (*11*). Microbial dysbiosis and immune dysfunction are therefore tightly intertwined, yet the causal direction of this relationship and their specific molecular drivers remain unresolved.

To dissect causality and address unresolved questions in the pathophysiology of CTLA-4 (haplo)insufficiency, we developed a physiologically relevant murine model system using a rewilding approach (*12, 13*), which has already significantly advanced disease modeling and translational research (*14*). In contrast to specific pathogen-free (SPF) counterparts, *CTLA4* heterozygous (*Ctla4⁺^/^⁻*) wildlings developed spontaneous skin disease, ocular secretions, diarrhea, and chronic colitis, mirroring the predominant manifestations observed in human *CTLA4* variant carriers. Longitudinal microbiome profiling demonstrated a progressive loss of α-diversity and an expansion of putative pathobionts, lagging clinical exacerbation -suggesting that host immune dysregulation and intestinal inflammation in turn promote sequential microbial disturbances. Moreover, immunophenotyping showed a sustained and pronounced skewing towards a pro-inflammatory immune milieu in the caecum of *Ctla4⁺^/^⁻* wildlings. Mechanistically, we further demonstrated that wildling microbiome-derived metabolites can drive inflammatory cytokine production in both human and murine T cells via activation of the NF-κB signaling pathway. These findings identify the microbiota as a critical modifier of disease penetrance in CTLA-4 haploinsufficiency and highlight *Ctla4⁺^/^⁻* wildlings as a prominent model to dissect host– microbiota interactions in immune dysregulation.

### The taxonomic and functional features of the gut microbiome are correlated with disease severity and clinical parameters

16S rRNA gene sequencing of stool from two independent cohorts previously revealed a correlation between the severity of CTLA-4 (haplo)insufficiency and the gut microbial composition (*10*). To achieve a more comprehensive characterization, we performed shotgun metagenomic sequencing of fecal microbiota in a larger newly assembled cohort with an expanded clinical metadata set. In total, 42 individuals carrying *CTLA4* variants were stratified into affected (high clinical score) and mildly affected or unaffected (low clinical score) groups, according to the CHAI scoring system for CTLA-4 (haplo)insufficiency (*15*) (Fig. 1A). In addition, healthy controls, including both related household members and unrelated non-household individuals, were included for comparative analysis. Consistent with previous 16S rRNA sequencing data (*10*), individuals with high clinical scores displayed markedly reduced microbial diversity compared with those with low clinical scores and to healthy controls (Fig. 1B). Microbial diversity was strongly correlated with clinical parameters (in norm = within reference range, low = below reference range, high = above reference range). Variant carriers with abnormal IgG or IgM levels, altered total leukocyte counts, reduced B cell counts, or decreased IgA levels exhibited significantly lower microbial diversity (Fig. 1C). To investigate compositional differences among the three groups further, we calculated the beta diversity using the Bray-Curtis (BC) distance, performed a permutational multivariate analysis of variance (PERMANOVA) controlled for confounders (sex, BMI, leukocyte number, B cell number, T cell number (Fig. S1A-B)), and confirmed a significant association between microbiome variation at the species level and disease severity (PERMANOVA, R2 = 0.0486 and p = 0.001; Fig. 1D, left). Similarly, we also observed that the disease severity was significantly associated with the microbiome functional composition (PERMANOVA, R2 = 0.052 and p = 0.001; Fig. 1D, right). Individual species analysis revealed a pronounced depletion of 37 bacterial taxa in the high clinical score group compared with healthy controls, including *Gemmiger formicilis* and *Lawsonibacter asaccharolyticus*, which are known carbohydrate-fermenting and butyrate-producing bacteria (*16, 17*). While no taxa were found to be enriched in the high clinical score group (Fig. 1E), the low clinical score group exhibited enrichment of 11 species relative to healthy controls (Fig. 1E).

**Fig. 1.**
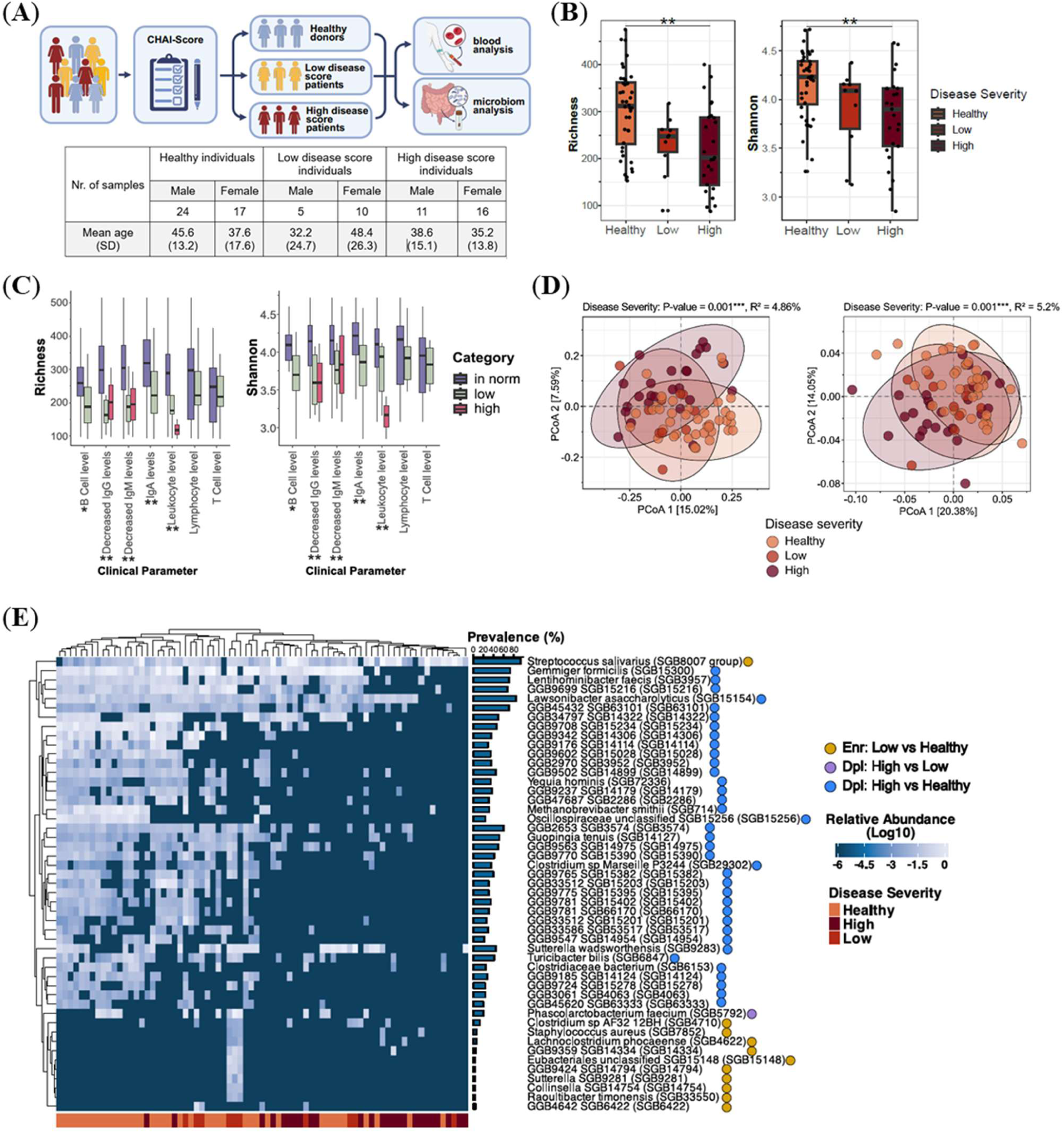
Shotgun metagenomic sequencing reveals microbial signatures associated with human CTLA-4 (haplo)insufficiency disease severity. A. Healthy controls (n = 41) and individuals with CTLA-4 (haplo)insufficiency (n = 42) were recruited from the Center for Chronic Immunodeficiency (CCI), University Medical Center Freiburg. *CTLA4* variant carriers were stratified into low-disease-score (Low, n = 15) and high-disease-score (High, n = 27) groups. PBMCs and clinical immunological parameters—including immunoglobulin levels and immune cell counts—were obtained from all participants. Stool samples from the same cohort were subjected to shotgun metagenomic sequencing. B. Alpha diversity (richness and Shannon index) is shown as box-and-whisker plots (median with interquartile range) across different clinical score groups. Statistical comparisons were performed with the Kruskal-Wallis test, followed by a post-hoc Dunn’s test, **p < 0.01. C. Alpha diversity measures (richness and Shannon) according to immune parameters (in norm = within reference range, low = below reference range, high = above reference range), shown as box plots with the line indicating the median and the whiskers representing the interquartile range. Statistical comparisons were performed with the Kruskal-Wallis test, followed by a post-hoc Dunn’s test, *p < 0.05, **p < 0.01. D. Beta diversity was assessed using PCoA based on Bray-Curtis dissimilarity matrices (see method). This analysis was performed for both species-level taxonomic profiles (left panel) and predicted functional pathways (right panel). The statistical significance of the bacterial community structure or functional pathway abundance was determined by PERMANOVA with 1,000 permutations based on the disease status. E. A heatmap illustrating significant differences in SGB-level bacterial abundances (enrichment or depletion) among healthy, low, and high groups. Yellow dots indicate species enriched in the low group compared with the healthy group; purple dots indicate species decreased in the high group compared with the low group; and blue dots indicate species decreased in the high group compared with the healthy group. Statistical significance was assessed using a one-tailed Wilcoxon rank-sum test, with FDR) correction applied using the Benjamini–Hochberg method (FDR < 0.05).

Analysis of specific microbial metabolic pathways by functional prediction revealed that the high clinical score group showed a marked loss of metabolic activity, including pathways involved in amino acid biosynthesis, branched-chain amino acid (BCAA) biosynthesis, glycogen biosynthesis, isoprenoid biosynthesis, fatty acid biosynthesis, the tricarboxylic acid (TCA) cycle, butyrate biosynthesis, myo-inositol degradation, and aminoacyl-tRNA biosynthesis. In contrast, fatty acid β-oxidation was elevated in this group (Fig. S1C). In the PCoA analysis, we observed significant differentiation of microbial taxa (Fig. S1D) and predicted functional profiles (Fig. S1E) between household and non-household healthy control communities. In addition, based on the abundance of shared species and predicted functional profiles, healthy donors living in the same household exhibited lower pairwise BC distances for both taxonomic (Fig. S1F) and functional (Fig. S1G) profiles, indicating greater compositional similarity within this group. Taken together, variant carriers with severe clinical symptoms and abnormal blood parameters demonstrate significantly lower microbial diversity, depletion of bacterial taxa, and impaired metabolic performance.

### *Ctla4^+/-^* wildlings spontaneously develop phenotypes resembling human CTLA-4 (haplo)insufficiency

To understand whether the microbiome is causally involved in the expressivity of the CTLA-4 (haplo)insufficiency phenotype, we generated *Ctla4⁺^/^⁻* wildlings by fostering SPF *Ctla4^+/-^* mice to wildling dams (*18*) to expose a naturalized microbiota from the day of birth (Fig. 2A). Both *Ctla4⁺^/^⁻* SPF and *Ctla4⁺^/^⁻* wildlings were monitored for 24 weeks, followed by comprehensive analysis including organ pathology, immune profiling, metabolomics, and longitudinal microbiome composition (Fig. 2A). Unlike the SPF counterparts which remained healthy, *Ctla4⁺^/^⁻* wildlings spontaneously recapitulated key features of human CTLA-4 (haplo)insufficiency. Although survival rates did not differ between groups (Fig. S2A), *Ctla4⁺^/^⁻* wildlings exhibited a significantly lower body weight from 5 weeks after birth onwards (Fig. 2B). As early as 2-3 weeks of age, *Ctla4⁺^/^⁻* wildlings developed ocular secretions and incomplete eyelid opening (Fig. 2C). Cutaneous symptoms followed a progressive course: subtle skin flaking appeared at 1–2 weeks (Fig. 2D), advancing to pronounced hair loss by 7 to 8 weeks (Fig. 2E, 2G middle). By week 5, diarrhea emerged (Fig. 2G left), consistent with intestinal inflammation as indicated by elevated stool lipocalin-2 concentrations (Fig. 2F). Longitudinal disease assessment revealed the trajectory of disease progression: disease severity peaked at 9–11 weeks and gradually subsided from week 13 onward (Fig. 2G). By week 24, a residual phenotype persisted, primarily in the colon and lung, manifesting as diarrhea, lymphocytic infiltration, and shortened colon (Fig. 2G-I, Fig. S2C). Another common clinical manifestation of human CTLA-4 (haplo)insufficiency is hypogammaglobulinemia. In our mouse model, we did not detect major differences in most immunoglobulin isotypes in the serum when comparing *Ctla4⁺^/^⁻*wildlings with SPF *Ctla4⁺^/^⁻*, except for elevated IgG1and IgE (Fig. S2B). This is partially attributable to the elevated baseline levels of IgG1, IgG2a, and IgE observed in wild-type mice following acquisition of a wild microbiota (Fig. S2B). Nevertheless, the reduced IgG2a levels in *Ctla4⁺^/^⁻* wildlings relative to wild-type wildlings suggest the presence of mild hypogammaglobulinemia at 24 weeks of age (Fig. 2J). In addition, we evaluated the spleen and mesenteric lymph node (mLN) weights across experimental groups. *Ctla4⁺^/^⁻* wildlings exhibited comparable absolute spleen and mLN weights, indicating no overt organ enlargement (Fig. 2K–L). However, both the spleen-to-body weight and mLN-to-body weight ratios were increased compared with wild-type wildlings at 24 weeks of age (Fig. S2D–E), indicating relative lymphoproliferation.

**Fig. 2.**
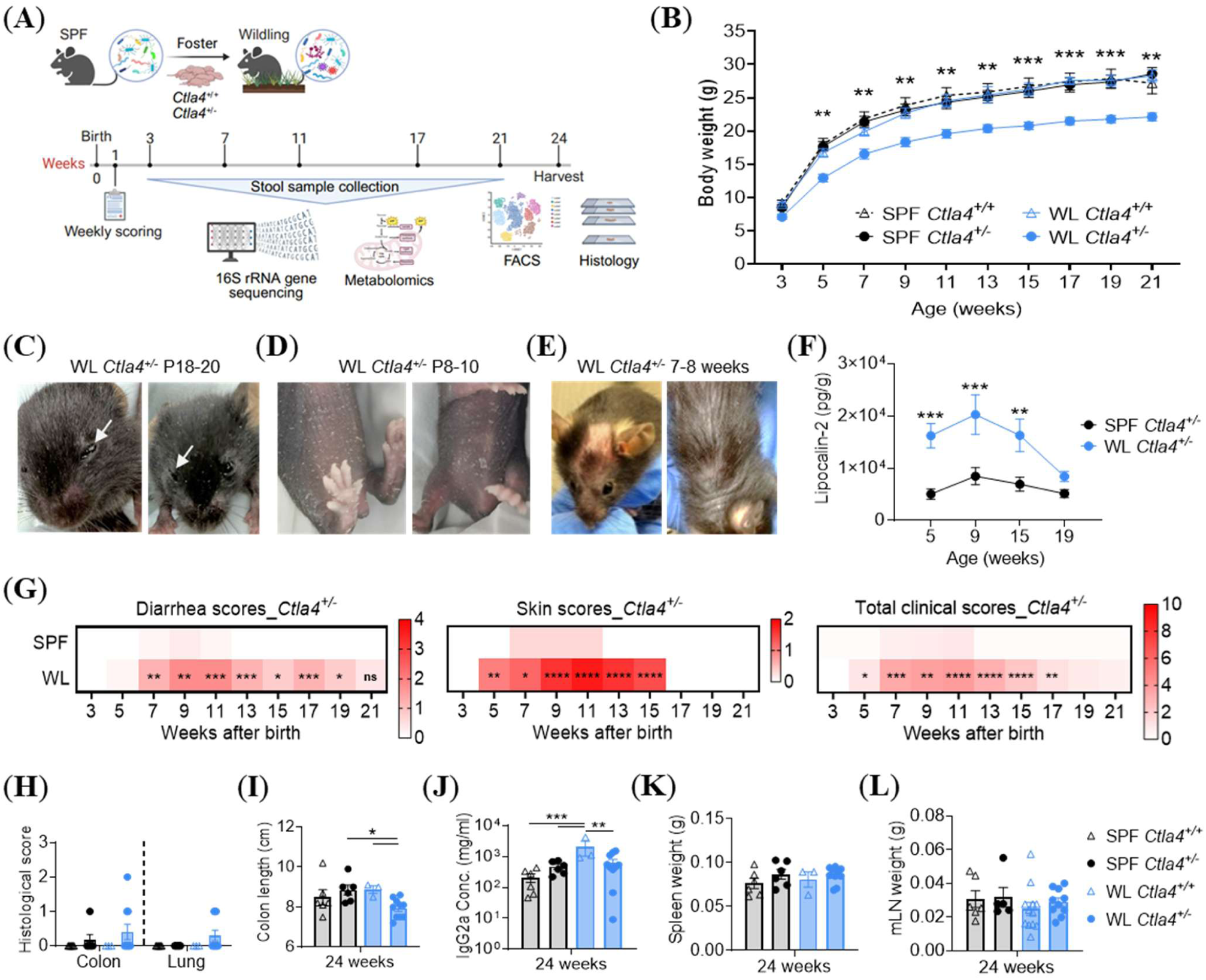
*Ctla4^+/-^* wildlings recapitulate the clinical phenotypes observed in humans with CTLA-4 insufficiency. A. Schematic overview of the experimental design, sampling time points, and analyses performed. B. Body weight of SPF and wildling (WL) *Ctla4^+/-^* and *Ctla4^+/+^* mice monitored from 3 to 21 weeks of age. * Indicates the comparison between SPF *Ctla4^+/-^* mice and *Ctla4^+/-^* wildlings. C-E. Representative images illustrating macroscopic disease manifestations (C: eye openness, D: skin flaking, E: hair loss) in *Ctla4^+/-^* mice colonized wildling microbiota. F. Stool lipocalin-2 concentrations measured by ELISA at 5, 9, 15, and 19 weeks of age in SPF and WL *Ctla4^+/-^* mice. G. Summary of the diarrhea, skin, and total clinical scores in SPF and WL *Ctla4^+/-^* mice from 3 to 21 weeks of age. Diarrhea scoring was based on stool consistency and anal cleanliness: 0 = normal stool; 1 = soft stool; 2 = soft stool with dirty anus; 4 = watery stool with dirty anus. Skin scoring was based on fur and skin integrity: 0 = normal (no fur loss, no skin lesions); 1 = fur loss; 2 = fur loss with skin rash. Total clinical scores were calculated as the sum of individual scores for body weight loss, diarrhea, bloody stool, skin condition, vocalization, eye openness, posture, activity, social behavior, and piloerection (see Table S1). Mice reaching a total score of 10 were considered to bear severe burden and were humanely euthanized. H. Hematoxylin and eosin (H&E) staining of colon and lung tissues derived from SPF and WL mice with corresponding histopathological evaluation. Scoring criteria for each organ are shown in Table S2. I. Colon length in SPF and WL *Ctla4^+/-^* and *Ctla4^+/+^* mice at 24 weeks of age. J. Serum IgG2a concentrations in SPF and WL mice with or without *Ctla4* mutation at 24 weeks of age. K-L. Spleen and mesenteric lymph nodes (mLNs) weights from SPF and WL mice at 24 weeks of age. Data in B, F, and G-J were obtained from two independent experiments and analyzed using a two-way ANOVA, n=3-10. *p < 0.05, **p < 0.01, ***p < 0.001, ****p < 0.0001.

In summary, *Ctla4⁺^/^⁻* wildlings recapitulate key clinical manifestations of human CTLA-4 (haplo)insufficiency, including impaired weight gain, ocular involvement, eczema, chronic intestinal inflammation, mild hypogammaglobulinemia and lymphoproliferation. Our findings indicate that exposure to a natural microbiota is indispensable for triggering CTLA-4 disease onset in case of (haplo)insufficiency. Moreover, the observed patterns of disease progression and remission highlight a dynamic co-adaptation among the microbiota, epithelial barrier integrity, and host immune responses, which collectively shape the longitudinal course of CTLA-4 (haplo)insufficiency.

### Longitudinal profiling of microbiome composition in SPF and wildling *Ctla4^+/-^* mice

To characterize the microbial development in *Ctla4^+/-^*mice under SPF and wildling condition, we longitudinally collected stool samples from individual mice and performed 16S rRNA gene sequencing. Principal coordinates analysis (PCoA) showed clear separation of microbial profiles between SPF and wildling groups, reflecting their distinct microbial compositions (Fig. 3A). Within the SPF group, the microbial community structure showed minor age-related variation. In contrast, *Ctla4⁺^/^⁻* wildlings exhibited pronounced, age-dependent shifts in the microbial composition over the study period (Fig. 3A). Alpha diversity analysis revealed a progressive decline in microbial richness in wildlings from weeks 3 to 17, reaching levels even lower than those of SPF counterparts (Fig. 3B). By comparison, microbial richness in SPF mice remained relatively stable across time (Fig. 3B). A similar trend was observed with the Shannon index indicating an overall reduction in species evenness in *Ctla4⁺^/^⁻* wildlings over time (Fig. 3B). This loss of diversity is consistent with observations in human *CTLA4* variant carriers, particularly those with high CHAI scores (Fig. 1B).

**Fig. 3.**
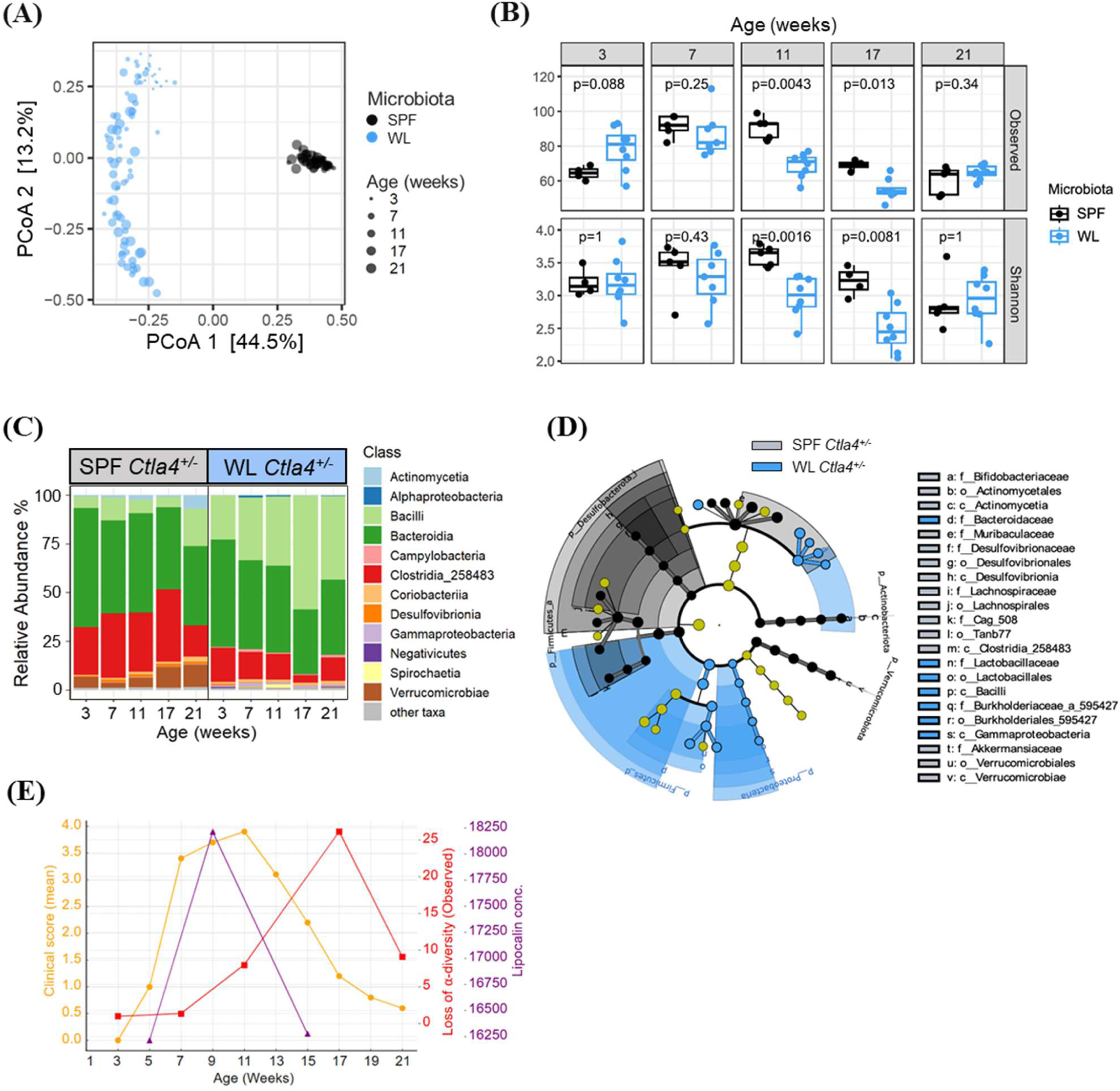
16S rRNA gene sequencing reveals dynamic microbial changes in SPF and WL *Ctla4^+/-^* mice between 3-21 weeks of age. Fecal pellets were collected longitudinally from individual mice between 3 and 21 weeks of age. DNA was extracted from the stool samples and subjected to 16S rRNA gene sequencing. A. â-diversity analysis (Principal Coordinate Analysis, PCoA) deconstructed distinct microbial community structures between SPF and WL mice, as well as age-associated shifts within individual mice over time. B. á-diversity metrics (observed features, top; Shannon index, bottom) for SPF and WL *Ctla4^+/-^*mice at five different ages. P-values were calculated using Welch’s t-test; n = 4– 8. C. Relative abundance of major bacterial classes in *Ctla4^+/-^* mice harboring SPF or WL microbiota at 5 different ages. D. Cladogram generated by LEfSe analysis showing bacterial taxa significantly enriched in fecal samples from SPF (gray) or WL (blue) *Ctla4*⁺^/^⁻ mice at 11 weeks of age. Concentric rings represent taxonomic levels from phylum (outer rings) to genus/species (inner nodes). Blue and black nodes and branches indicate taxa with significantly different relative abundances between the two groups (Kruskal–Wallis test, *p* < 0.05), whereas yellow nodes indicate taxa without significant differences. E. Mean clinical score (orange, left y-axis), mean loss of activity (red, right y-axis), and mean lipocalin concentration (purple, far right y-axis) from the same cohort of animals are shown as a function of age (weeks).

We next compared the microbial taxonomic profiles of *Ctla4^+/-^*mice housed under wildling or SPF conditions. Across weeks 3 to 21, Actinomycetia, Bacteroidia, Clostridia_258483, and Verrucomicrobiae were enriched in SPF mice, whereas Alphaproteobacteria, Bacilli, Campylobacteria, Gammaproteobacteria, Negativicutes, and Spirochaetia were preferentially enriched in wildlings (Fig. 3C). At 11 weeks of age, the time point corresponding to the peak clinical score, LEfSe analysis further identified significant increases in Gammaproteobacteria and Burkholderiaceae in the wildling group (Fig. 3D), taxa that have been implicated in intestinal inflammation and are frequently associated with inflammatory bowel disease (IBD) (*19, 20*). In contrast, Akkermansia, a genus well known for its role in strengthening mucosal barrier integrity (*21*), was significantly enriched in SPF mice (Fig. 3D). To determine the impact of the genotype on the microbial composition in wildlings, we compared wildling *Ctla4^+/+^* and *Ctla4^+/-^* littermates. At the phylum level, the communities were indistinguishable—except for the emergence of Spirochaetota in *Ctla4^+/-^* mice at the age of week 11 (Fig. S3A), when the clinical score of the mice peaked. At week 11, none of the *Ctla4^+/+^* animals harbored Spirochaetota, whereas three of eight *Ctla4^+/-^* mice were positive (Fig. S3B). These three mice also had markedly elevated lipocalin-2 levels at week 9 when compared with the Spirochaetota-negative *Ctla4^+/-^*mice (Fig. S3C). These findings suggest a possible association between Spirochaetota and severe gastrointestinal pathology in our mice. When aligning clinical scores, lipocalin-2 levels, and microbial diversity loss by age, we observed that microbial dysregulation lagged behind disease onset by approximately 6 weeks (Fig. 3E), suggesting that host immune factors, such as the *Ctla4* mutation, contribute to subsequent microbial dysregulation.

### Microbial metabolites in the intestinal compartment of *Ctla4^+/-^* wildlings exhibit a pro-inflammatory profile

In addition to characterizing the gut microbial composition, we performed untargeted metabolomic profiling of fecal samples collected across the various disease stages. At week 12, corresponding to the peak of clinical disease severity, principal component analysis (PCA) demonstrated a clear microbiota-dependent separation of fecal metabolic profiles (Fig. 4A). This divergence persisted through week 22 (Fig. S4A). Hierarchical clustering consistently showed that metabolite profiles segregated primarily by microbiota rather than by age (Fig. 4B).

**Fig. 4.**
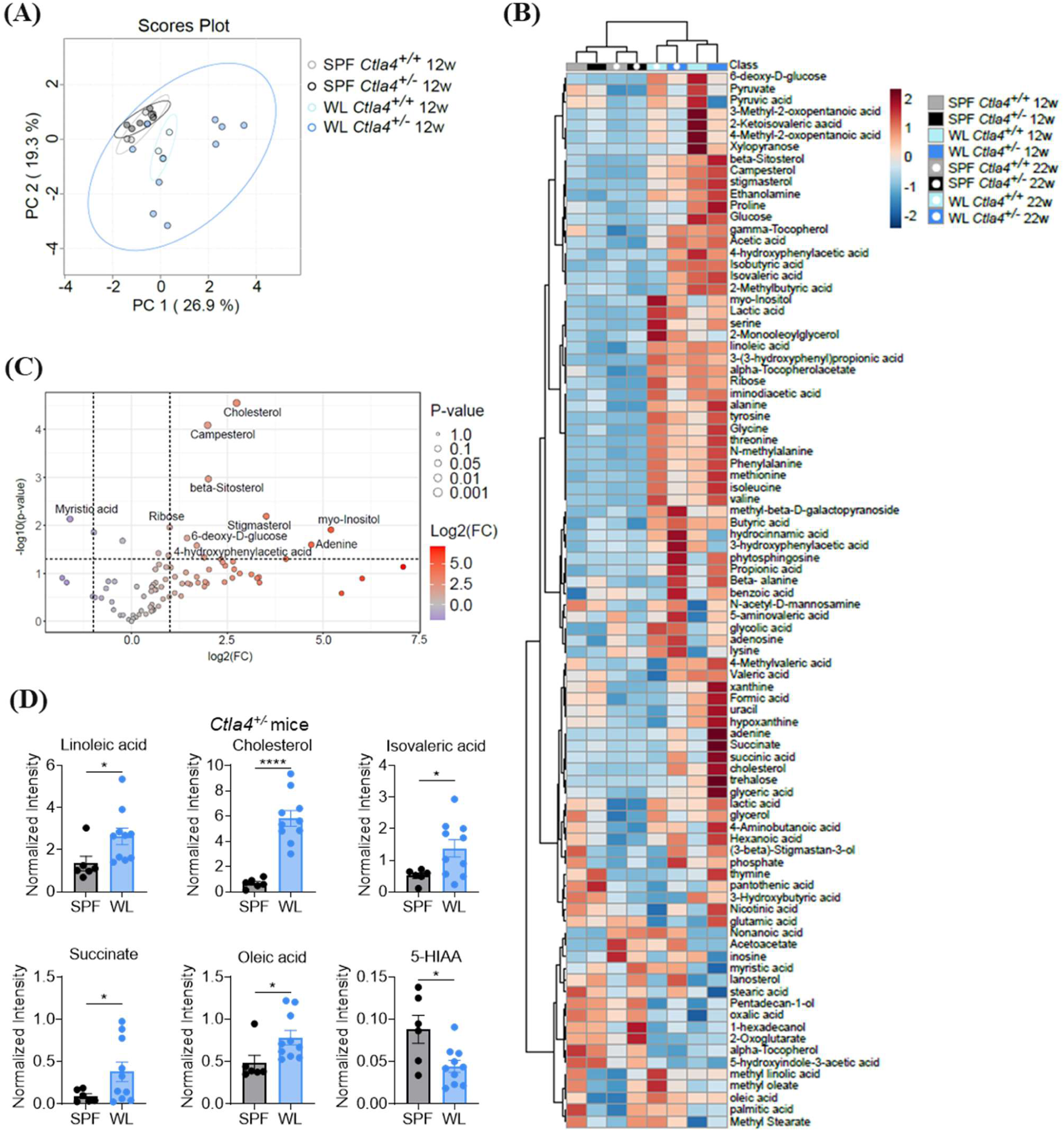
Using untargeted metabolic measurements to characterize metabolic changes in the gut when colonizing wildling microbiota. A. Principal component analysis of fecal metabolites shows a clear differentiation between 12-week-old SPF and wildlings. n=3-10. Ellipses show 95 % confidence. B. Heat map displaying z-scores of averaged metabolite intensities per group. Groups and metabolites are clustered. n=3-10. C. Volcano plot comparing fecal metabolites of 12-week-old *Ctla4^+/-^* wildlings with *Ctla4^+/-^* SPF mice with log2 foldchange and negative decadic logarithm of the p-values. Dashed lines indicate a foldchange of 0.5 or 2 as well as a p-value of 0.05. n=6-10 D. Bar charts of metabolites with immunomodulatory properties in 12-week-old wildling and SPF *Ctla4^+/-^* mice. Data were obtained from two independent experiments and analyzed using Welch’s t test, n=6-10. *p < 0.05, ****p < 0.0001.

At week 12, *Ctla4⁺^/^⁻* wildlings displayed a broad metabolic reprogramming characterized by a significant enrichment of multiple pathways, including amino acid metabolism (e.g. valine, glycine, threonine, serine, tyrosine, isovaleric acid, isobutyric acid, 4-hydroxyphenylacetic acid), carbohydrate and central energy metabolism (e.g. glucose, ribose, 6-deoxy-D-glucose, succinic acid, glyceric acid), short-chain fatty acids (SCFAs) (e.g. acetate, propionate), lipid metabolism (e.g. cholesterol, campesterol, β-sitosterol, linoleic acid, oleic acid), and ethanolamine metabolism (Fig. 4B–C). Several metabolites elevated in *Ctla4⁺^/^⁻* wildlings such as linoleic acid, cholesterol, isovaleric acid, succinate, and oleic acid (Fig. 4D) have been implicated in NF-κB activation, reactive oxygen species (ROS) signaling, and pro-inflammatory cytokine production (*22–24*). Conversely, levels of 5-hydroxyindoleacetic acid (5-HIAA), a microbiota-regulated metabolite that signals through the aryl hydrocarbon receptor (AhR), were markedly reduced in *Ctla4⁺^/^⁻* wildlings (Fig. 4D). Ahr signaling promotes Treg differentiation, stability, and function, particularly at mucosal surfaces such as the gut, and has documented protective effects in experimental models of colitis and arthritis (*25–27*),

By week 22, wildlings maintained a metabolic profile largely consistent with week 12, with additional enrichment of alanine, isoleucine, 3-hydroxyphenylacetic acid (3-HPAA), and phenylalanine, whereas SPF mice continued to exhibit elevated levels of 5-HIAA (Fig. 4B, S4B). Notably, SPF mice showed no major metabolic shifts between weeks 12 and 22 (Fig. S4C). In *Ctla4⁺^/^⁻* wildlings, several metabolites including thymine, glutamic acid, uracil, glyceric acid, formic acid, and β-sitosterol were significantly higher at week 12, whereas nonanoic acid, stearic acid, acetoacetate, and pyruvic acid were preferentially enriched at week 22 (Fig. 4B).

### *Ctla4^+/-^* wildlings exhibit a distinct immune state and T cell phenotype

Microbial metabolites play a critical role in immune cell development, maturation, and functional differentiation. We therefore sought to determine how the wildling microbiome and its associated metabolic profile influence host immune homeostasis. To comprehensively characterize immune phenotypes, we performed 28-color spectral flow cytometry on samples from the spleen, mesenteric lymph nodes (mLN), and cecum tissue using a Sony spectral analyzer. The panel was designed to capture major lymphoid and myeloid compartments, including T cell subsets, B cells, innate immune populations, and innate lymphoid cells (ILCs) (Fig. S5A). Colonization with a wildling microbiota substantially reshaped the baseline immune composition, as evidenced by the distinct immune profile of *Ctla4^+/+^* wildlings compared with SPF *Ctla4^+/+^* controls (Fig. 5A). Next, we evaluated the impact of the *Ctla4* mutation on the immune profile. Comparison of SPF *Ctla4^+/+^* and SPF *Ctla4^+/-^* mice revealed only modest immune perturbations, including a slight increase in Th1 frequencies in both spleen and mLN (Fig. 5A). Strikingly, *Ctla4⁺^/^⁻* wildlings exhibited a markedly altered immune landscape relative not only to SPF *Ctla4^+/-^*mice but also to their *Ctla4^+/+^* wildling counterparts (Fig. 5A). This included significantly reduced NK cells and B cells, accompanied by elevated frequencies of ILC3s, γδ T cells, T follicular helper (Tfh) cells, and exhausted CD8^+^ T cells across all three tissues examined (spleen, mLN, and cecum) (Fig. 5A). These observations indicate that the *Ctla4* mutation reshapes the immune landscape only in the presence of a natural microbiota, but not under SPF conditions.

**Fig. 5.**
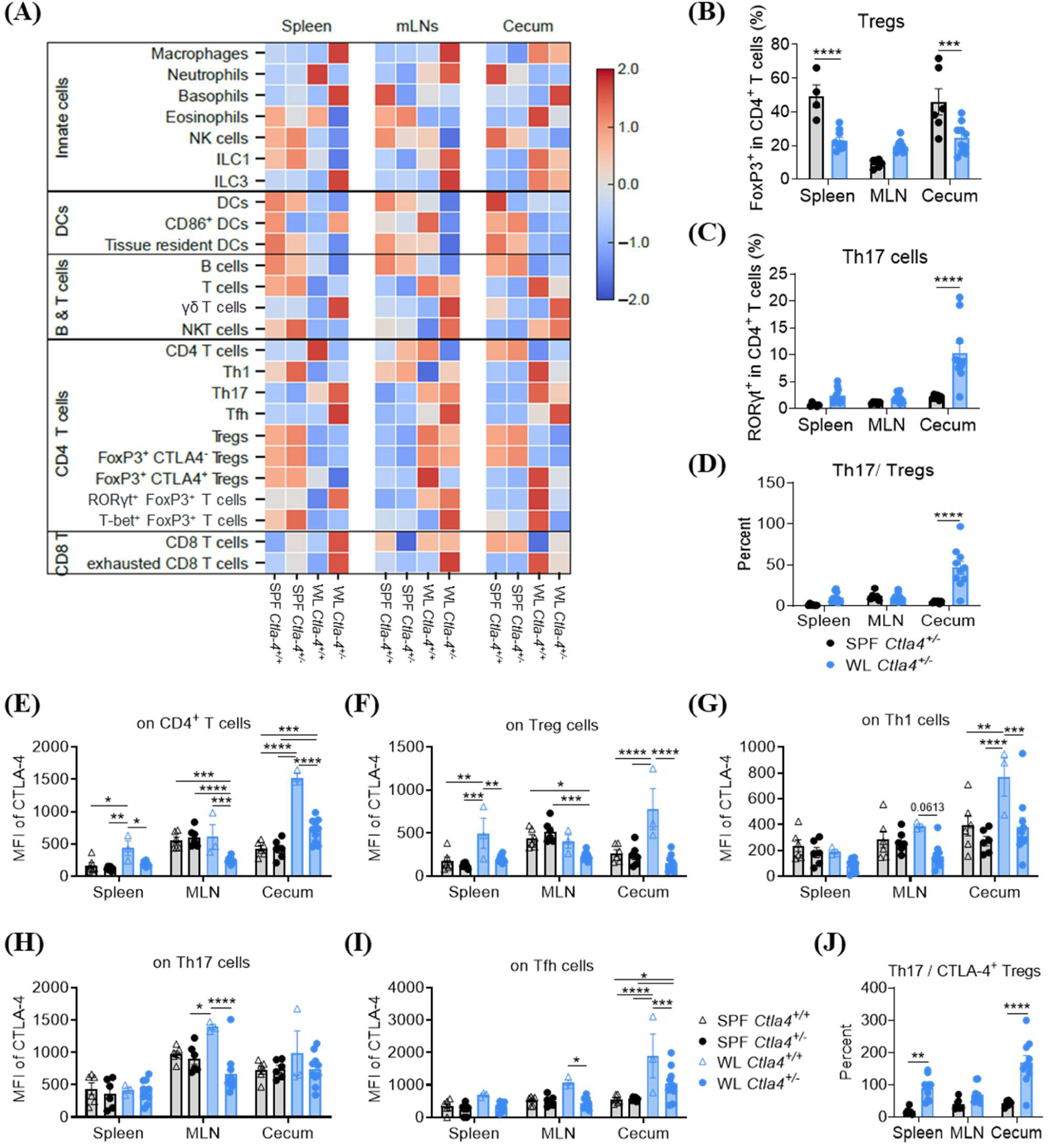
Features and functions of innate and adaptive immune cells in secondary lymphoid organs and intestine of SPF and wildlings (WL). Lymphocytes from the spleen, mesenteric lymph nodes (mLNs), and cecum of SPF or WL mice at age of 24 weeks were isolated and stained with a 28-color antibody panel and analyzed using a Sony ID7000 spectral cytometer. A. Heatmap depicting the immune landscape across four groups and three organs. Each row represents the z-score–normalized frequency of an immune cell subset relative to its parental population. The color scale reflects standardized expression levels, with red indicating higher-than-average and blue indicating lower-than-average abundance within each cell type, n=3-10. B. Frequencies of FoxP3^+^ cells within the CD4^+^ T cell compartment in the spleen, mLN, and cecum of SPF or WL *Ctla4^+/-^*mice, n=4-10. C. Frequencies of RORγt^+^ cells within the CD4^+^ T cell compartment of SPF or WL *Ctla4^+/-^* mice, n=6-10. D. Ratios of Th17 cells to Tregs, calculated by dividing the frequency of Th17 cells by the frequency of Tregs within each individual mouse, n=6-10. E-I. Mean fluorescence intensity (MFI) of CTLA-4 in total CD4^+^ T cells (E), FoxP3^+^ CD4^+^ T cells (F), Tbet^+^ CD4^+^ T cells (G), RORγt^+^ CD4^+^ T cells (H), and ICOS^+^ CXCR5^+^ CD4^+^ T cells (I) across the 3 organs of SPF or WL mice, n=3-10. J. Ratios of Th17 cells to CTLA-4^+^ Tregs, calculated by dividing the frequency of Th17 cells by the frequency of CTLA-4^+^ Tregs within each individual mouse, n=6-10. Data in B-J represent two independent experiments and were analyzed by two-way ANOVA. *p < 0.05, **p < 0.01, ***p < 0.001, ****p < 0.0001.

Given the central role of CTLA-4 in regulating T cell activation and tolerance, we next focused on T cell subset frequencies and phenotypes. Th17 and Treg cells represent key pro-inflammatory and regulatory T cell populations in the intestine, respectively. *Ctla4^+/-^* mice housed under wildling conditions exhibited significantly reduced Treg frequencies (Fig. 5B) and elevated Th17 frequencies (Fig. 5C) compared with SPF *Ctla4^+/-^* mice across all tissues, resulting in an increased Th17/Treg ratio, most prominently in the cecum (Fig. 5D). These data suggest that the wildling microbiota promotes a more pro-inflammatory intestinal environment, which may be particularly detrimental in the context of CTLA-4 (haplo)insufficiency. We further evaluated the CTLA-4 protein levels across T cell subsets. Under SPF conditions, CTLA-4 mean fluorescence intensity (MFI) on total CD4⁺ T cells was comparable between *Ctla4^+/+^* and *Ctla4^+/-^* mice in all organs (Fig. 5E). In contrast, *Ctla4^+/+^* wildlings exhibited significantly increased CTLA-4 level, particularly within the cecum. Remarkably, *Ctla4⁺^/^⁻* wildlings expressed only approximately half the CTLA-4 levels on CD4⁺ T cells compared with wild-type controls (Fig. 5E). Analysis across Treg, Th1, Th17, and Tfh subsets revealed consistently reduced CTLA-4 level in *Ctla4⁺^/^⁻*wildlings than the wildling wild-type counterparts, although the magnitude of reduction varied by subset (Fig. 5F-I). Moreover, we sought to validate T cell phenotypes and functional states by analyzing gene expression in colon tissue samples. Although not statistically significant, we observed a trend toward decreased *Il10* expression, increased *Il17* expression, and decreased *Ctla4* expression at the mRNA level in *Ctla4⁺^/^⁻* wildlings compared with *Ctla4^+/+^*wildlings (Fig. S5B). In summary, both the wildling microbiota and *Ctla4* mutation exert broad and synergistic effects on innate and adaptive immune development. *Ctla4⁺^/^⁻* wildlings exhibit long-term impaired T cell regulation and compromised immune tolerance when challenged with a natural microbial environment.

### Microbiome-derived metabolites from *Ctla4^+/-^* wildlings directly induce proinflammatory cytokine secretion in mouse and human T cells

Given the pronounced differences in gut metabolomes and immune landscapes between SPF and wildlings, we wondered whether the microbiome-derived metabolites directly influence T cell inflammatory programming. To answer this question, we performed *ex vivo* stimulation assays using fecal “stool water” on primary mouse and human cells. To obtain fecal stool water, fecal pellets from wildling and SPF *Ctla4^+/-^* mice were weighed and homogenized in PBS (1:10 w/v). After centrifugation, supernatants were collected and passed through a 0.2-µm filter to remove viable microbes. These preparations were applied on splenic naïve T cells (Fig. S6A) sorted from SPF *Ctla4^+/-^* mice for 16 hours, after which culture supernatants were harvested for cytokine quantification (Fig. 6A). Stool water derived from *Ctla4⁺^/^⁻* wildlings, but not from SPF *Ctla4^+/-^* mice, robustly induced IL-1α secretion from naïve mouse T cells in a dose-dependent manner (Fig. 6B). Other cytokines were either unchanged or showed no clear response pattern following stool water stimulation (Fig. S6B). Notably, this response was not attributable to inflammatory cytokines present in the stool water itself: both SPF and wildling stool water samples used for stimulation exhibited low or undetectable levels of IL-23, IL-1α, IFN-γ, TNF-α, MCP-1, IL-12p70, IL-6, IL-17A, and GM-CSF (Fig. S6C).

**Fig. 6.**
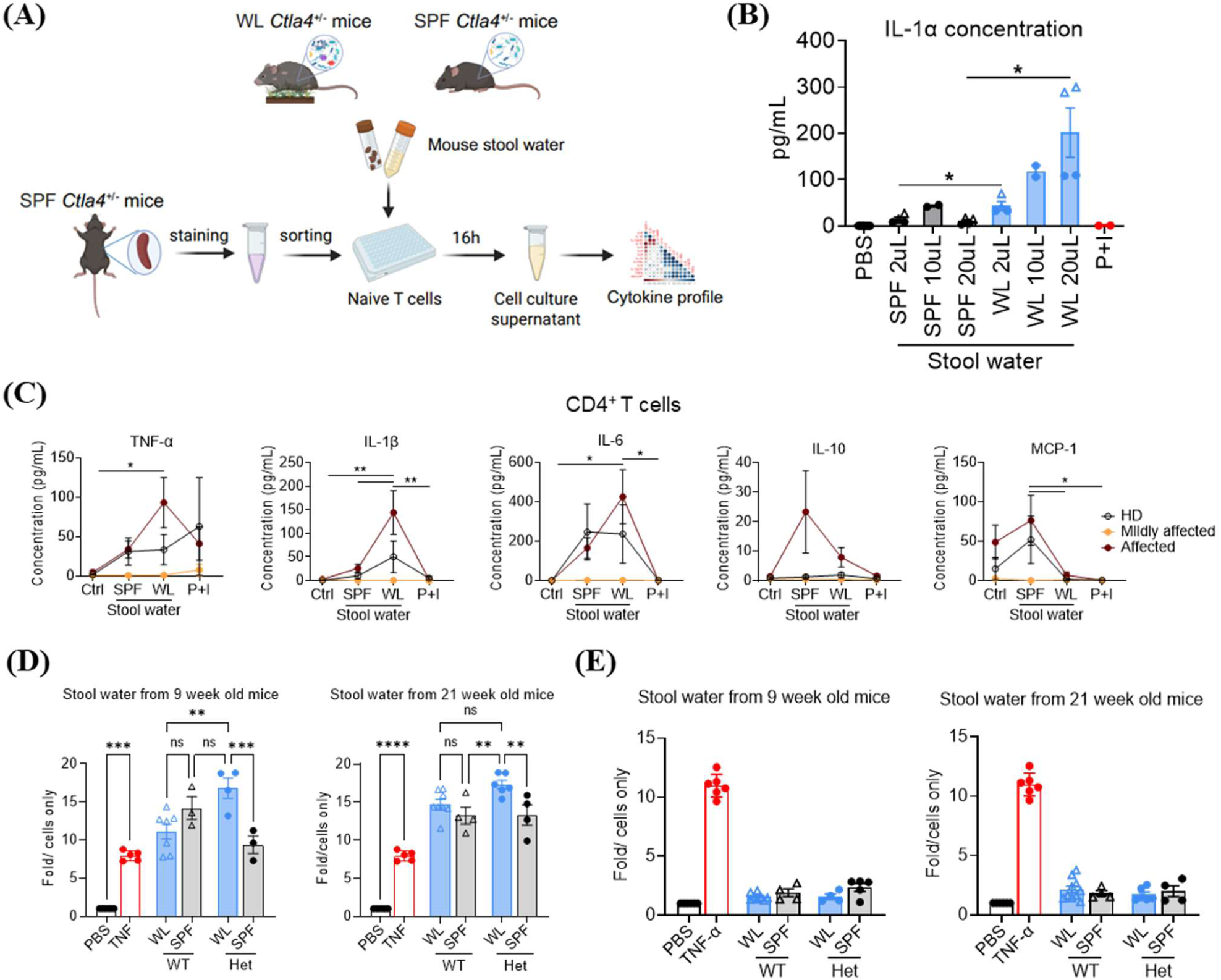
*Ex vivo* stimulation of T cells with stool water from SPF and WL mice. A. Schematic overview of the experimental setup. Fecal pellets collected from SPF or WL mice were resuspended in PBS at a 1:10 (w/v) ratio and thoroughly homogenized. After centrifugation, the supernatant was collected and filtered to remove microbial cells. B. Naïve splenic CD4⁺ T cells derived from SPF *Ctla4^+/-^*mice were incubated for 16 hours with PBS, varying amounts of SPF or WL stool water from 11-week-old mice, and PMA+ionomycin (P+I). Culture supernatants were collected, and IL-1α concentrations were measured by ELISA. Circles and triangles represent responses induced by stool water from two different mice within each group. P-values were calculated using Welch’s t-test, *p < 0.05, n=2-8. C. Naïve CD4⁺ T cells isolated from human PBMCs were incubated for 16 hours with PBS, SPF or WL stool water (from 11-week-old mice), and P+I. Culture supernatants were collected, and the concentrations of multiple cytokines were analyzed using a LEGENDplex assay. Data represent two independent experiments and were analyzed by two-way ANOVA. *p < 0.05, **p < 0.01, n=2-6. D-E. Jurkat NF-κB luciferase reporter cells (D) or LS174T NF-κB luciferase reporter cells (E) were incubated for 4 hours with PBS (vehicle control), mouse stool water derived from 9- or 21-week-old *Ctla4^+/+^* wildling, *Ctla4^+/–^* wildling, *Ctla4^+/+^* SPF, *Ctla4^+/–^* SPF mice, or TNF-α (positive control). NF-κB activation was quantified by luciferase activity. Data are shown as fold induction relative to PBS-treated (cells only) control and are presented as mean ± SD. Each dot represents stool water derived from an individual mouse. Data were analyzed by one-way ANOVA. **p < 0.01, ***p < 0.001, ****p < 0.0001, n=3-10.

We next applied the same stimulation strategy to naïve CD4⁺ T cells isolated from PBMCs (Fig. S6D) of individuals with CTLA-4 (haplo)insufficiency. Remarkably, wildling stool water induced strong production of inflammatory cytokines including TNF-α, IL-1β, and IL-6 especially in T cells from clinically affected *CTLA4* variant carriers, whereas healthy donors or mildly affected individuals exhibited moderate or low cytokine induction, respectively (Fig. 6C). In contrast, SPF stool water induced IL-10 production in naïve T cells from affected *CTLA4* variant carriers, and MCP-1 production in cells from affected individuals and healthy donors (Fig. 6C). Together, these results demonstrate that wildling microbiota–derived metabolites directly modulate T cell function, driving inflammatory cytokine secretion in both mouse and human naïve T cells. This effect is particularly pronounced in individuals with CTLA-4 (haplo)insufficiency who exhibit more severe immune dysregulation, suggesting that microbial metabolic cues synergize with impaired CTLA-4 signaling to exacerbate inflammatory T cell responses. Given that NF-κB is a central regulator of pro-inflammatory cytokine expression, including IL-1β, IL-6, and TNF-α (*28,29*), we next assessed whether stool water derived from our mouse cohorts exhibited differential NF-κB–activating capacity. To this end, we performed *in vitro* stimulation assays using two NF-κB reporter cell lines: Jurkat T cells and a human colorectal epithelial cell line LS174T. Stool water induced markedly stronger NF-κB activation in Jurkat T cells than in epithelial cells (Fig. 6D–E). Notably, stool water from *Ctla4^+/–^* wildlings elicited significantly higher NF-κB activity compared with samples from *Ctla4^+/+^*wildlings and *Ctla4^+/–^* SPF mice (Fig. 6D, left). This enhanced activation was evident in samples from 9-week-old mice, corresponding to peak disease severity. At 21 weeks, *Ctla4^+/–^* wildling-derived stool water remained significantly more stimulatory than *Ctla4^+/–^* SPF samples, but not *Ctla4^+/+^* wildling samples (Fig. 6D, right). In contrast, epithelial reporter cells exhibited minimal differential activation across groups suggesting microbiota-derived metabolites, antigens, or molecules from *Ctla4^+/–^*wildlings preferentially induced inflammatory cytokine production through activation of the NF-κB signaling pathway, particularly in CTLA-4 insufficient T cells. Together with our primary T cell stimulation data, these results support a model in which microbiota-derived signals amplify inflammatory T cell responses via enhanced NF-κB pathway activation, thereby providing a mechanistic link between microbial dysbiosis, aberrant T cell activation, and disease heterogeneity in CTLA-4 haploinsufficiency.

## Discussion

In this study, we established a wildling *Ctla4^+/-^* mouse model that developed spontaneous clinical features closely mirroring those observed in humans with CTLA-4 (haplo)insufficiency. By integrating longitudinal clinical assessment, microbiota profiling, high-dimensional immune phenotyping, and metabolomic analyses, we investigated the causal role of the microbiome in disease pathogenesis in CTLA-4 (haplo)insufficiency. Our findings reveal that (i) a natural microbiota acts as a potent environmental trigger that drives disease manifestation and induces profound alterations in both innate and adaptive immune compartments, and (ii) immune dysregulation and intestinal inflammation in turn promote secondary microbial disturbances. Moreover, *ex vivo* functional assays demonstrate that microbiota-derived metabolites promote inflammatory cytokine production in both murine and human naïve CD4 T cells via NF-κB signaling pathway. Differential responsiveness of primary human T cells from *CTLA4* variant carriers to microbiome-derived metabolites may contribute to clinical heterogeneity. These insights offer a mechanistic explanation for the variable penetrance seen in CTLA-4 (haplo)insufficiency and support microbiome- or metabolite-targeting strategies as potential therapeutic approaches.

Due to the asymptomatic phenotype of SPF *Ctla4^+/–^* mice, approaches to generate CTLA-4 (haplo)insufficiency mouse models for dissecting the disease pathophysiology have been developed, such as RNA-mediated knockdown of *Ctla4* expression (*30, 31*) and inducible deletion strategies (*32*). However, the clinical relevance of these models to human CTLA-4 (haplo)insufficiency is limited. Here, we present a complementary model based on microbiome manipulation that recapitulates key features of the human phenotype and enables direct investigation of the microbiota’s disease-modifying effects. Notably, prior work has shown that WildR mice also develop colitis following immune checkpoint blockade (ICB) (*33*), further underscoring the relevance of natural microbial exposure in CTLA-4–related pathology. Consistent with this concept, both human and mouse studies have attempted to mitigate irAEs through fecal microbiota transplantation (FMT) or administration of defined bacterial consortia (*34–36*), highlighting the potential of microbiome-based therapeutics in CTLA-4 insufficiency. However, our longitudinal study in *Ctla4^+/–^* wildlings suggests that these beneficial effects of microbiome-based interventions may be transient, potentially due to persistent host immune dysfunction driving subsequent microbial dysregulation.

Beyond CTLA-4 (haplo)insufficiency, the wildling platform offers powerful opportunities to study other inborn errors of immunity (IEIs). Our previous work demonstrated that wildling *gp91^-/-^* and *p47^-/-^* mice faithfully phenocopy human chronic granulomatous disease (CGD), an IEI caused by variants in components of the NADPH oxidase complex (*8*). In that model, we identified specific pathobionts—such as *Corynebacterium kutscheri*—that directly drove disease manifestations, mirroring observations in patients. Together, these observations highlight that wildlings represent a valuable system for studying IEIs that do not manifest under SPF conditions and emphasize the urgency of delineating genetic predisposition–microbiome interactions across different IEIs. Moreover, wildlings may serve as an advanced preclinical model for testing interventions targeting the microbiota–host–immune interface.

The microbiome is well known to shape the abundance and functionality of regulatory T (Treg) cells in the gut (*37–40*). Our findings extend this concept by suggesting that CTLA-4 expression on Tregs, and across multiple T helper cell subsets, is itself strongly influenced by microbial exposure. Under SPF conditions, basal CTLA-4 expression appears sufficient to maintain immune homeostasis, whereas in wildling wild-type animals, increased microbial complexity drives robust CTLA-4 upregulation in Tregs and other T cells, providing critical immunosuppressive control. Consistent with our findings, recent studies have demonstrated that CTLA-4 expression in innate lymphoid cells (ILCs) is also regulated by the microbiota, where intestinal colonization and microbiota-induced IL-23 signaling promote CTLA-4 expression on ILC3s (*41, 42*). Increased CTLA-4 expression has also been reported in ILC1s from patients with colitis. How wild microbiota in our model elicits CTLA-4 expression in T cells remains to be fully elucidated. CTLA-4 expression in Tregs is known to be induced by T-cell receptor (TCR) signaling, activation, and proliferation (*43*). We therefore propose that strong TCR engagement—particularly by diverse microbial antigens present in wildling environments—promotes Treg activation and CTLA-4 upregulation to restrain excessive immune responses. Another study demonstrated that Tregs can acquire lactate, which promotes CTLA-4 expression through an RNA-splicing– mediated mechanism (*44*). Consistent with this observation, we detected a marked enrichment of lactate in the gut metabolome of wildlings. These findings raise the possibility that lactate, or other microbiota-derived metabolites, may contribute to the increased CTLA-4 expression. However, this microbiome-mediated CTLA-4 upregulation appears to be substantially impaired in *Ctla4⁺^/^⁻* wildlings, resulting in a pathogenic immune environment and insufficient immune suppression at the intestinal barrier.

Microbial metabolites are heavily involved in / regulating immune cell by shaping cellular energy metabolism, engaging host receptors, and modulating signaling pathways that influence cytokine production (*45*). In our study, untargeted metabolomic analyses revealed that the gut metabolite profile was primarily shaped by the wild microbiota rather than host genotype. These wildling microbiome-derived metabolites, molecules or antigens were sufficient to directly activate naïve murine T cells *ex vivo*, and trigger robust production of TNF-α, IL-1β, and IL-6 in T cells from clinically affected human *CTLA4* variant carriers via activation of NF-κB signaling pathway. Many metabolites-including amino acids, α-ketoglutarate, lactate, saturated fatty acids (SFAs), and succinate-have been shown to activate NF-κB signaling (*46, 47*). Our next approach will be to target these metabolites in order to mitigate exacerbated NF-κB activation and the associated production of inflammatory cytokines. Interestingly, divergent responses of human PBMCs to wildling microbiome–derived metabolites were observed. Cells from mildly affected individuals showed minimal activation, highlighting the profound influence of gene–environment interactions on disease penetrance and severity and providing a potential mechanistic explanation for the observed disease heterogeneity.

One notable phenotype of *Ctla4⁺^/^⁻*wildlings is the gradual attenuation of disease severity from approximately 15 weeks of age onward, suggesting that additional host compensatory mechanisms may partially restore microbial and immune homeostasis. Previous studies have reported that regulatory T cells (Tregs) isolated from human PBMCs exhibit compensatory upregulation of immunosuppressive cytokines (e.g., IL-10) and activation markers such as GITR, ICOS, and OX40 during anti-CTLA-4 monoclonal antibody treatment (*48*). Other immune cell populations may also contribute to this compensatory process. Indeed, prior studies have shown that CTLA-4 blockade or Treg-specific CTLA-4 deficiency can induce PD-1 upregulation in CD8⁺ T cells, promoting the expansion of exhausted CD8⁺ T cells (*49, 50*). A detailed investigation of Treg transcriptomic profiles, as well as the functional states of other adaptive immune cell populations in *Ctla4⁺^/^⁻* wildlings, will therefore be required to clarify the mechanisms underlying this compensatory response. Moreover, whether this recovery *Ctla4^+/-^* wildlings is transient or sustained, and whether disease relapse may occur with aging, remains unknown. Our future work will involve extended longitudinal monitoring and immune profiling at multiple disease stages to resolve these questions.

Despite the strength of our findings, it is worth noticing that all *Ctla4*^+/-^ mice in this study share the same targeted exon knockout, which does not encompass the genetic heterogeneity seen in human CTLA-4 (haplo)insufficiency. Patients often carry missense variants that disrupt CTLA-4 folding (*51*), surface trafficking (*52*), or ligand binding (*52*), resulting in diverse biochemical outcomes, possibly additionally influencing the clinical presentation. Thus, although our model recapitulates key disease features, additional models incorporating patient-specific alleles will be required to fully capture the spectrum of CTLA-4 dysfunction.

Nonetheless, our integrated mouse–human analyses identify the microbiome—and particularly its metabolic outputs—as a central determinant of disease penetrance in CTLA-4 (haplo)insufficiency. By demonstrating that natural microbial exposure and microbiota-derived metabolites directly shape T cell responses, we provide mechanistic insight into how environmental factors modulate monogenic immune disorders. These findings highlight therapeutic opportunities in targeting microbial communities or metabolic pathways to ameliorate disease in CTLA-4 insufficiency and potentially irAEs and other IEIs.

## Acknowledgments

We thank all patients and healthy donors for their generous participation and for providing blood and stool samples. We thank the Lighthouse Core Facility (LCF) for their support with cell sorting, FACS, etc. We thank Pavla Mrovecova for laboratory organization, coordination between clinical and laboratory teams, and technical support. The authors used AI-assisted tools exclusively for linguistic editing and grammar correction. Figures 1A, 2A, and 6A were created using BioRender.

## Author contributions

Conceptualization: BZ, BG

Investigation: BZ, LS, PS, KDH, SL, TRL, WS, AH, KG, CS, VA

Visualization: BZ, LS, PS, SL, TRL, WS, AH, VA

Funding acquisition: BG, SPR

Project administration: BZ, BG

Supervision: BZ, BG, SPR, BK, TS

Writing – original draft: BZ

Writing – review & editing: BG, TS, SL, PS, KDH, VA, LS, RG

## Data and materials availability

All sequencing data generated in this study will be deposited in a public repository prior to publication. All other data supporting the findings of this study are available within the paper or the supplementary materials. Materials generated in this study are available from the corresponding author upon reasonable request.

