## Supplementary Materials for "The microbiome determines the phenotype in CTLA-4 insufficient mice and men"

**This PDF file includes:**

Materials and Methods
Figures S1 to S6
Tables S1 and S2

### Materials and Methods

Human Shotgun Metagenomic Sequencing and Data Processing

Raw metagenomic sequencing reads underwent quality control and preprocessing using the BBMap suite (specifically the BBDuk tool; sourceforge.net/projects/bbmap/). Low-quality bases were trimmed, and reads mapping to the human host genome or phiX contaminants were removed to ensure non-host data purity *(53)*.

- Taxonomic and functional profiling

Taxonomic profiling of the microbial communities was performed on the pre-processed reads using MetaPhlAn 4 *(54)* with default parameters. Taxa were filtered based on abundance; only bacteria present at a relative abundance greater than 0.001 (0.1%) at the Species-level Genome Bin (SGB) level were retained for downstream analysis.

Functional profiling was conducted using the HUMAnN3 pipeline *(55)*. The resulting pathway abundance tables were normalized to relative abundance to account for sequencing depth differences across samples.

- Assessment of community diversity and principal coordinates analysis (PCoA)

Community diversity metrics were calculated using the vegan package in R. Alpha-diversity was assessed via community richness and the Shannon diversity index. For beta-diversity analysis, relative abundance data were first transformed using an arcsine square-root transformation to stabilize variance. Subsequently, Bray-Curtis dissimilarity matrices were calculated and visualized using Principal Coordinate Analysis (PCoA). Statistical significance of the bacterial community or functional pathway was evaluated using Permutational Multivariate Analysis of Variance (PERMANOVA) with 1,000 permutations.

- Bacterial enrichment/depletion abundance

Bacterial enrichment/depletion analysis was performed using pairwise comparisons across disease severity groups (i.e., Healthy vs. Low Severity, High vs. Low Severity, and Healthy vs. High Severity). Differences in bacterial abundance were assessed using a one-tailed Wilcoxon Rank-Sum Test to identify taxa significantly enriched in specific groups. P-values were adjusted for multiple testing using the Benjamini-Hochberg (BH) False Discovery Rate (FDR) method.

- Household vs. non-household comparison of bacterial and functional composition

To assess household versus non-household microbial similarity, Bray-Curtis distances for both taxonomic and functional profiles were calculated between samples from the same household and from different households. Statistical comparisons were performed using two-tailed Wilcoxon rank-sum tests.

Mice

SPF *Ctla4*$^{+/-}$ mice on C57BL/6J background were purchased from Jackson Laboratory (B6.129P2Ctla4<tm1Mak>/Mmjax). Wildling C57BL/6NTac mice were created by embryo transfer of laboratory mouse into pseudopregnant wild mice and provided by Stephan Rosshart lab *(56)*. All mice were housed and bred at the Medical Center – University of Freiburg, Germany. Throughout the entire experiment, mice were handled under BSL1 conditions and were housed under a 12:12 light:dark cycle (room temperature 20–24°C, humidity 45%-65%, air exchange rate of 15 times per hour) in a Green Line IVC system from Tecniplast inside of autoclaved microisolator cages (GM500) with autoclaved rodent chow (GRANOVIT AG, KLIBA NAFAG, 3437.PX.L15) and autoclaved tap water ad libitum, 1x autoclaved sizzle pad 8 g (ssniff), 1x autoclaved play tunnel (ssniff), and Aspen wood chip bedding (ssniff). Experiments were performed in accordance with the guidelines of the Federation for Laboratory Animal Science Associations (Felasa) and the national animal welfare body. They were in compliance with the German animal protection law and were approved by the animal welfare committee of the Regierungspräsidium Freiburg (permit G-21/151, and G-25/050). Animals used in experiments were gender and age matched. All mice were euthanized by asphyxiation with $CO_2$.

Genotyping

Ear-punch tissue collected at week 3 was lysed in Ear-Punch Buffer with 400 µg/mL Proteinase K and incubated overnight at 56°C. Proteinase K was heat-inactivated at 95°C for 10 minutes, after which 250 µL of $H_2O$ was added to each lysate and the tubes were vortexed thoroughly.
From this crude DNA preparation, 1 µL was used in a 20 µL PCR reaction. PCR amplification was performed using the cycling conditions listed below, followed by agarose gel electrophoresis to resolve the PCR products.

| Name | Sequence |
|---|---|
| f-Mut-Ctla4 | ATT-TGT-CAC-GTC-CTG-CAC-GAC-G |
| f-Wt-Ctla4 | CCC-CAC-AAT-TGG-AAA-CTC-TC |
| r-Com-Ctla4 | ACC-CTA-TCC-TTC-CTG-CTC-CT |

| Step | Temp °C | Duration | Note |
|---|---|---|---|
| 1 | 94.0 | 2 min | |
| 2 | 94.0 | 30s | |
| 3 | 65.0 | 30s | - 0.5°C per cycle decrease |
| 4 | 68.0 | 30s | |
| 5 |  | ---- | repeat step 2-4 for 10 cycles (Touchdown) |
| 6 | 94.0 | 30s | |
| 7 | 60.0 | 30s | |
| 8 | 72.0 | 30s | |
| 9 | | --- | repeat step 6-8 for 28 cycles |
| 10 | 72.0 | 1 min | |
| 11 | 10.0 | hold | |

Fostering

Timed pregnant breeding pairs of wildling wild-type (WT) and SPF *Ctla4*$^{+/-}$ x *Ctla4*$^{+/+}$ were set up. The SPF pups were transferred to their wildling foster mothers immediately after birth, and tail clipping was used to distinguish the pups. Wildlings were kept under semi-natural housing conditions through cage supplementation with natural materials, including hay, compost, and fomites from actual wild mice.

16S rRNA gene amplicon sequencing

Fecal pellets were collected from all mice at the indicated time points, snap frozen, and stored at −80°C. First, genomic DNA was extracted using the ZymoBIOMICS 96 MagBead DNA Kit according to the manufacturer's instructions. 16S rRNA gene amplification of the V4 region

(F515/R806) was performed according to an established protocol previously described *(58)*. Briefly, DNA was normalized to 25 ng/µl and used for sequencing PCR with unique 12-base Golay barcodes incorporated via specific primers (obtained from Sigma). PCR was performed using Q5 polymerase (NewEnglandBiolabs) in triplicates for each sample, using PCR conditions of initial denaturation for 30 s at 98°C, followed by 25 cycles (10 s at 98°C, 20 s at 55°C, and 20 s at 72°C). After pooling and normalization to 10 nM, PCR amplicons were sequenced on an Illumina MiSeq platform via 250 bp paired-end sequencing (PE250). Resulting raw reads were demultiplexed by idmp (https://github.com/yhwu/idemp) according to the given barcodes *(57)*. Libraries were processed including merging the paired end reads, filtering the low-quality sequences, dereplication to find unique sequences, singleton removal, denoising, and chimera checking using the USEARCH pipeline version 11.0.667 *(58)*. In brief, reads merged by fastq_mergepairs command (parameters: maxdiffs 30, pctid 70, minmergelen 200, maxmergelen 400), filtered for low quality with fastq_filter (maxee 1) and singletons using fastx_uniques command (minuniquesize 2). We run UPARSE algorithm to make 97% OTUs and filter chimeras calling, following the amplicon quantification using usearch_global command (strand plus, id 0.97, maxaccepts 10, top_hit_only, maxrejects 250). Taxonomic assignment was conducted by Constax (classifiers: rdp, sintax, blast) using the GreenGenes2 database *(59)* and summarizing into biom-file for the visualization and downstream analysis in phyloseq in the R statistical programming *(60)*. To determine bacterial OTUs that explained differences between microbiota settings, the LEfSe method was used. OTUs with Kruskal-Wallis test <0.05 and LDA scores >3.5 were considered informative.

<u>Extraction and analysis of fecal metabolites</u>

Metabolites were extracted from wet feces samples on ice with 1 ml extraction solution (acetonitrile:methanol 3:1, v:v, containing internal standards). After centrifugation (4°C, 20,000 g, 10 min), extracts were removed and the remaining feces samples were dried in a vacuum concentrator. The extracts were diluted with the extraction solution to equal dry weight concentrations. 300 µl, corresponding to 1 mg dry feces, were evaporated in a new reaction tube in a vacuum concentrator for untargeted GC/MS based metabolic profiling. 100 µl were used for targeted SCFA analysis.

GC/MS analysis was conducted as previously reported *(61)*: Metabolite pellets were derivatized by methoxyamine hydrochloride and N-methyl-N-trimethylsilyl trifluoroacetamide, separated on an HP-5MS column and analyzed by a mass spectrometer equipped with an EI-source. Runs were aligned and features identified by different spectral libraries based on spectral match and retention index.

SCFA analysis was adapted from Tan *et al.*: Carboxylic acids were derivatized by 1-Ethyl-3-(3-dimethylaminopropyl) carbodiimide and O-benzyl-hydroxyl amine in pyridine buffer *(62)*. After liquid-liquid extraction with chloroform, analytes were separated on an eclipse C18 column coupled to a triple-quadrupole mass spectrometer.

All intensities were normalized to an internal standard and range scaled. Statistical analysis and visualization were done with MetaboAnalyst 6.0 *(63)*.

<u>Cell Isolation from Spleen, mLN, and cecum</u>

The spleen and mLN from each animal were placed on a 70 µm cell strainer placed on top of a 50 mL Falcon tube and mechanically dissociated using a syringe plunger. The strainer was then washed with PBS to a final volume of 10 mL for spleen samples and 5 mL for mLN samples.

Aliquots of 10 µL spleen or 10 µL mLN were taken for cell counting. All samples were centrifuged at 1600 rpm for 10 minutes at 4°C, after which the supernatant was discarded.
The cecum was cleaned of fat and opened longitudinally. The tissue was washed twice in PBS, then incubated for 20 minutes in 10 mL dissociation buffer (HBSS without MgCl2 (Gibco) with 2 mM EDTA (Sigma)+ 10 mM HEPES (Sigma)) at 37°C with gentle shaking (130 rpm). Following incubation, the tissues were vortexed, cut into small pieces and transferred into a 50 mL Falcon tube containing 5 mL digestion buffer (DMEM with 2% FCS, 10 mg/mL DNase I, and 0.5 mg/mL Collagenase Type IV). Samples were incubated for 20 minutes at 37°C on a shaker at 130 rpm. After incubation, samples were shaken vigorously 10 times and passed through a 70 µm strainer. The remaining tissue was mechanically dissociated using a syringe plunger and washed 1 time with 10 mL DMEM + 5% FCS. The lamina propria (LP) cell suspension was centrifuged at 1600 rpm for 10 minutes at 4°C. The cell pellet was resuspended in 4 mL of 40% Percoll, and this suspension was carefully layered on top of the 80% Percoll. The gradient was centrifuged at 450 × g for 25 minutes at room temperature with acceleration setting 2 and deceleration setting 0. Lymphocytes were collected from the interphase and transferred to a new 15 mL Falcon tube.

Flow cytometry

Spleen, mLN, and cecum cells were transferred into a U-bottom 96-well plate, 100 µL of Fc-blocking mix was added to each well, mixed by gentle pipetting, and incubated for 5 minutes at 4°C. Cells were then washed with 100 µL FACS buffer and centrifuged at 1600 rpm for 5 minutes at 4°C. For surface staining, 50 µL of the surface antibody mixture as listed below was added to each well, and cells were incubated for 20 minutes at 4°C in the dark. Cells were washed with 200 µL FACS buffer and centrifuged at 1600 rpm for 5 minutes at 4°C. Cells were then fixed by resuspending them in 200 µL fixation buffer and incubating for 30 minutes at 4°C in the dark. After fixation, cells were washed with 100 µL Perm buffer at 2100 rpm for 5 minutes at 4°C, followed by an additional wash with 250 µL Perm buffer under the same conditions. For intracellular staining, the intracellular/nuclear antibody mixture as listed below was added, and cells were incubated overnight at 4°C. Cells were washed with 200 µL FACS buffer at 2100 rpm for 5 minutes at 4°C, resuspended in 200 µL FACS buffer for acquisition on the SONY ID7000 cytometer.

| Surface Marker | Fluorochrome | Titration |
|---|---|---|
| CD19 | BUV395 | 1:1000 |
| live/dead | LiveDeadFixableBlue | 1:1000 |
| TCRγδ | BUV496 | 1:100 |
| ST2 | BUV563 | 1:1000 |
| NKP46 | BUV661 | 1:250 |
| Gr-1/ Ly-6G | BUV737 | 1:400 |
| CD185/ CXCR5 | BV421 | 1:400 |
| FcεR1α | PacBlue | 1:400 |
| CD3 | BV510 | 1:400 |
| CD45 | BV570 | 1:400 |
| PD1 | BV605 | 1:100 |
| Siglec-F | BV650 | 1:250 |

| | | |
|---|---|---|
| CD278/ICOS | BV711 | 1:1000 |
| CD11b | FITC | 1:1000 |
| CD4 | NovaFluorBlue660 | 1:400 |
| F4/80 | PerCP | 1:400 |
| CD86 | PE-Dazzle594 | 1:100 |
| IL-7R (CD127) | PE-Cy5 | 1:250 |
| NK1.1 | PE-Fire700 | 1:1000 |
| CD196/CCR6 | PE-Fire810 | 1:400 |
| CD152/ CTLA-4 | APC | 1:400 |
| CD11c | SparkNIR-685 | 1:400 |
| CD8 | APC-H7 | 1:400 |

| Intracellular/ Intranuclear Marker | Fluorochrome | Titration |
|---|---|---|
| RORγt | BV785 | 1:100 |
| Gata3 | BB700 | 1:400 |
| T-bet | PE | 1:100 |
| IFN-γ | PE-Cy7 | 1:250 |
| FoxP3 | AF700 | 1:100 |
| CD152/ CTLA-4 | APC | 1:400 |

RNA isolation and quantitative PCR
RNA from colon tissue was isolated using the ExtractMe Total RNA Kit (Cat# EM09.2, Blirt), followed by concentration measurement with a NanoDrop 2000 spectrophotometer (Thermo Scientific). 1 mg of total RNA was used to generate cDNA using RevertAid First Strand cDNA Synthesis Kit (Cat# K1622, Thermo Scientific). Real-Time PCR was performed using gene-specific primer sets of *Gapdh*, *Il10*, *Il17*, *Ctla4*, and Kapa Sybr Fast qPCR kit (Applied Biosystems) on StepOnePlus Real-time PCR system (Applied Biosystems). PCR conditions were 95°C for 60 s, followed by 40 cycles of 95°C for 3 s and 60°C for 30 s. Data were analyzed using the deltaCt method with Gapdh serving as the reference housekeeping gene.

Histology
All organ samples were collected, fixed in 4% paraformaldehyde (PFA) and embedded in paraffin according to standard histological procedures. Sections of 2 µm thickness were stained with hematoxylin-eosin (HE) and evaluated by light microscopy blinded to the experimental groups.

Immunoglobulin Quantification
Blood was collected by cardiac puncture into and centrifuged after clotting (3000 × g, 10mins, 20 °C). Serum was collected and stored at −80 °C until further use. For antibody isotyping, a Mouse Immunoglobulin Isotyping Panel (6-plex) (Cat# 740493) LEGENDplex kit from BioLegend and Mouse IgE ELISA MAX™ Standard Set kit (Cat# 432401) from BioLegend were used according to the manufacturer's instructions.

Samples were diluted 1:50,000 to measure the immunoglobulin subtypes IgG1, IgG2a, IgG2b, IgG3, IgA, and IgM. Sample dilution for IgE measurement was adjusted according to microbial composition: SPF mice 1:10, wildlings 1:500.
Data acquisition of Immunoglobulin measured by LEGENDplex kit was performed on a BD LSRFortessa equipped with three lasers 405 nm, 488 nm, 640 nm. IgE was analyzed by the Tecan Spark plate reader at OD450nm and OD600nm.

*Ex vivo* incubation and cytokine measurement

To obtain fecal stool water, fecal pellets from *Ctla4*$^{+/-}$ wildlings or SPF *Ctla4*$^{+/-}$ mice were weighed and homogenized in PBS (1:10 w/v) *(64)*. After centrifugation (21000g, 10mins, 4°C), supernatants were collected and passed through a 0.2-µm filter to remove viable microbes. These preparations were applied on splenic naïve T cells sorted from SPF *Ctla4*$^{+/-}$ mice for 16 hours, after which culture supernatants were harvested for cytokine quantification using LEGENDplex Human inflammation Panel 1 (13-plex) (Cat# 740809) or LEGENDplex Mouse Inflammation Panel (13-plex) (Cat# 740446). Data acquisition of the cytokines measured by LEGENDplex kit was performed on a BD LSRFortessa equipped with three lasers 405 nm, 488 nm, 640 nm.

NF-κB reporter assay

NF-κB activation was assessed using Jurkat T cell and LS174T human colon epithelial cell NF-κB luciferase reporter lines, as previously described1. Cells were maintained in complete RPMI (Jurkat) or DMEM (LS174T) supplemented with 10% fetal bovine serum and antibiotics under standard culture conditions (37°C, 5% $CO_2$).
Reporter cells were plated in 96-well plates and stimulated with PBS (vehicle control), filtered stool water (10% v/v), or recombinant murine TNF-α (positive control) for 4 hours. Following stimulation, cells were lysed and NF-κB driven luciferase expression was assessed using the Pierce™ Firefly Luc One-Step Glow Assay Kit (ThermoFisher Scientific) according to the manufacturer's instructions. NF-κB activation was expressed as fold induction relative to PBS-treated control wells (cells only). Each biological replicate represents stool water derived from an individual mouse *(64)*.

Lipocalin-2 measurement

The stool samples collected across all experimental groups were weighed and homogenized in PBS (1:10 w/v). After centrifugation (21000g, 10mins, 4°C), supernatants were collected and lipocalin-2 concentrations were measured using the Mouse Lipocalin-2/NGAL Quantikine ELISA Kit from R&D systems (Cat# MLCN20) according to manufacturer's instructions. Absorbance was measured at 450 nm and 570 nm using a Tecan Spark plate reader.

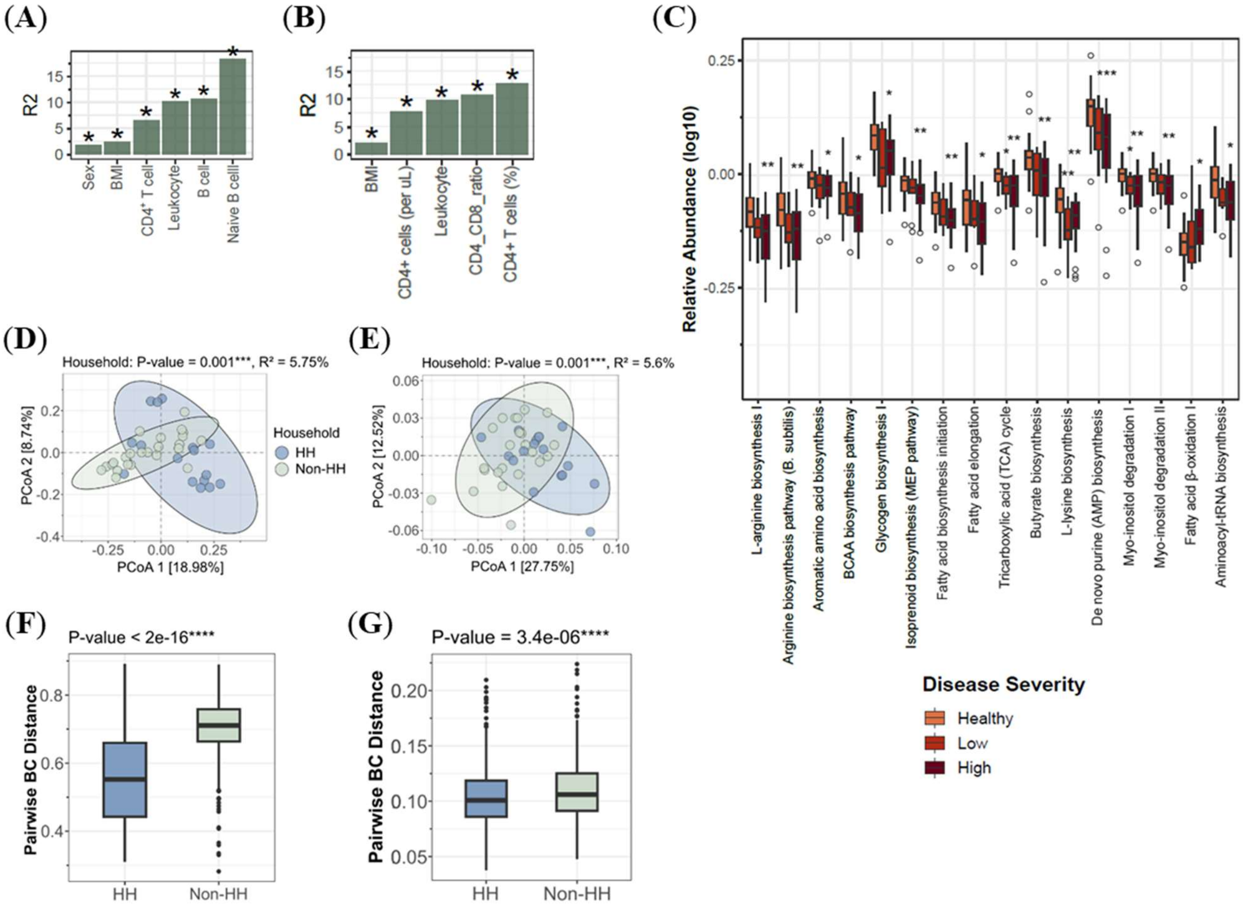


**Figure S1. Human CTLA-4 (haplo)insufficiency cohort metagenomics sequencing analysis.**

A. Covariance analysis of disease severity status and clinical parameters assessed at the SGB level of the gut bacterial community using PERMANOVA (1,000 permutations). The R2 values indicate the percentage of variance in bacterial composition explained by each clinical parameter. Significance is denoted by *p < 0.05.

B. Covariance analysis of disease severity status and clinical parameters assessed at the functional pathway level of the gut bacterial community using PERMANOVA (1,000 permutations). The R2 values indicate the percentage of variance in functional pathway composition explained by each clinical parameter. Significance is denoted by *p < 0.05.

C. Comparison of dominant functional pathways using the Kruskal-Wallis test, followed by a post-hoc Dunn's test. P-values were adjusted for multiple comparisons using the Benjamini-Hochberg (BH) false discovery rate (FDR) method. Significance levels are denoted as: *FDR < 0.05, **FDR < 0.01, ***FDR < 0.001.

D. PCoA of gut bacterial composition at the SGB level, illustrating clustering patterns among intra-household and inter-household individuals. Dissimilarity was calculated using Bray-

Curtis distance and statistical significance was assessed via PERMANOVA (1,000 permutations).

E. PCoA of gut microbiome functional pathways, illustrating clustering patterns among household and non-household individuals. Dissimilarity was calculated using Bray-Curtis distance and statistical significance was assessed via PERMANOVA (1,000 permutations).
F. Pairwise comparison of bacterial dissimilarity (Bray-Curtis distance at the SGB level) between household and non-household pairs. Significance was determined using two-tailed Wilcoxon rank-sum tests.
G. Pairwise comparison of functional dissimilarity (Bray-Curtis distance at the functional pathway level) between intra-household and inter-household pairs. Significance was determined using two-tailed Wilcoxon rank-sum tests.

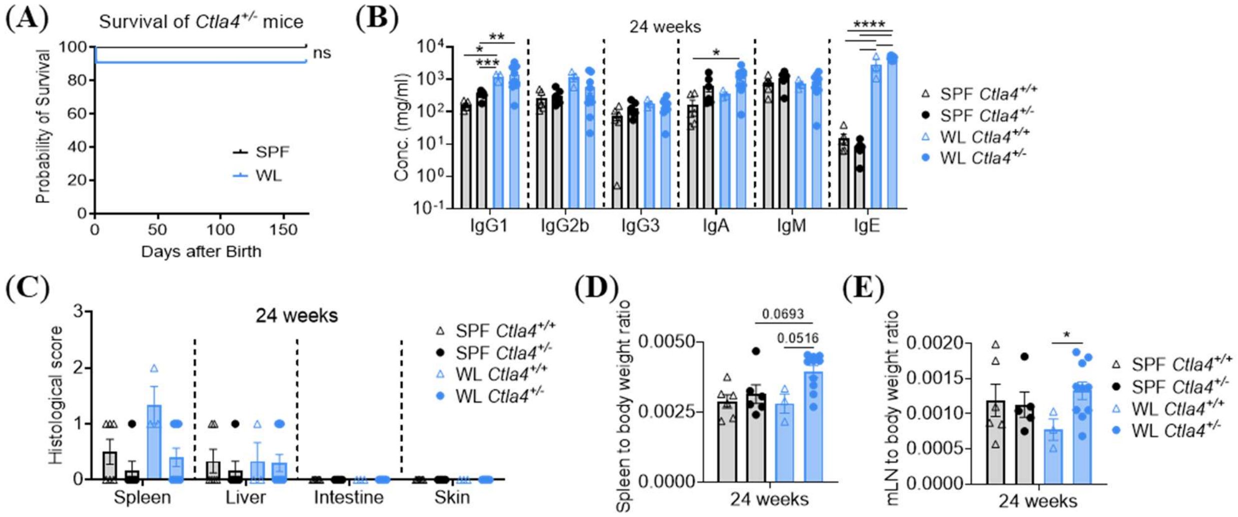


**Figure S2. Subtle discrepancies in survival, immunoglobulin isotypes, organ pathology, spleen weight, and mLN weight between WL *Ctla4*$^{+/-}$ mice and SPF *Ctla4*$^{+/-}$ mice at 24 weeks of age.**

A. 24-week survival rate of *Ctla4*$^{+/-}$ mice SPF or WL microbiome conditions. Survival curves were analyzed using Log-rank (Mantel-Cox) test, n=6-11. ns: non-significant.

B. Serum immunoglobulin isotype levels in SPF and WL mice with or without *Ctla4* mutation at 24 weeks of age. Statistical comparisons were performed using two-way ANOVA, n=3-10, *$p < 0.05$, **$p < 0.01$, ***$p < 0.001$, ****$p < 0.0001$.

C. H&E staining of spleen, liver, intestine, and skin tissues derived from SPF and WL mice at 24 weeks of age accompanied by corresponding histopathological evaluation. Scoring criteria for each organ are provided in Table 2.

D-E. Spleen weight to body weight ratio (D) and mLN to body weight ratio (E) of SPF or WL mice at 24 weeks of age. Data obtained from two independent experiments and analyzed using a two-way ANOVA, n=3-10. *$p < 0.05$.

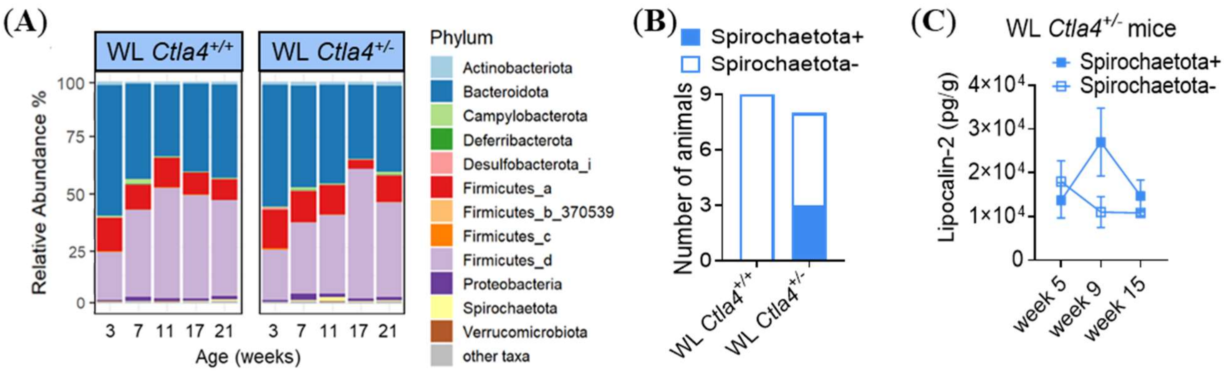


**Figure S3. The microbial comparison between WL *Ctla4*$^{+/-}$ and WL *Ctla4*$^{+/+}$ mice.**

A. Relative abundance of major bacterial phyla in WL *Ctla4*$^{+/-}$ and WL *Ctla4*$^{+/+}$ mice at 5 different ages. n=8-9

B. Numbers of Spirochaetota positive and negative animals in WL *Ctla4*$^{+/-}$ and WL *Ctla4*$^{+/+}$ group at 11 weeks. n=8-9

C. Stool lipocalin-2 concentrations measured by ELISA at 5, 9, and 15 weeks of age in WL *Ctla4*$^{+/-}$ mice. n=3

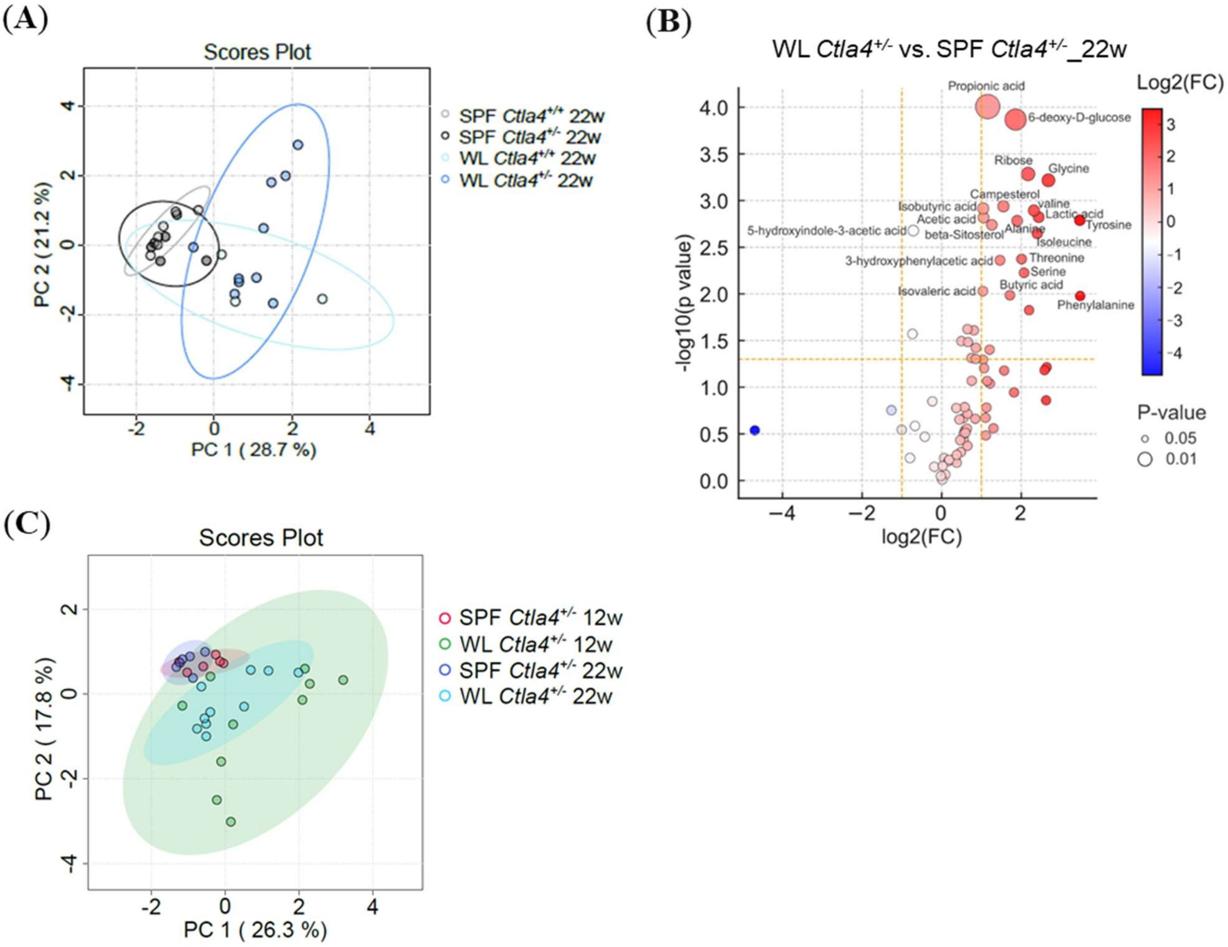


**Figure S4. Fecal metabolites in wildlings and SPF mice.**

A. Principal component analysis (PCA) of fecal metabolites shows a clear differentiation between 22-week-old SPF and wildlings (WL). n=3-10. Ellipses show 95 % confidence.
B. Volcano plot comparing fecal metabolites of 22 weeks old *Ctla4*$^{+/-}$ wildlings with *Ctla4*$^{+/-}$ SPF mice with log2 fold change and negative decadic logarithm of the p-values. Dashed lines indicate a foldchange of 0.5 or 2 as well as a p-value of 0.05. n=6-10
C. PCA of fecal metabolites confirmed age-independent separation between wildling and SPF mice. n=6-10. Ellipses show 95 % confidence.

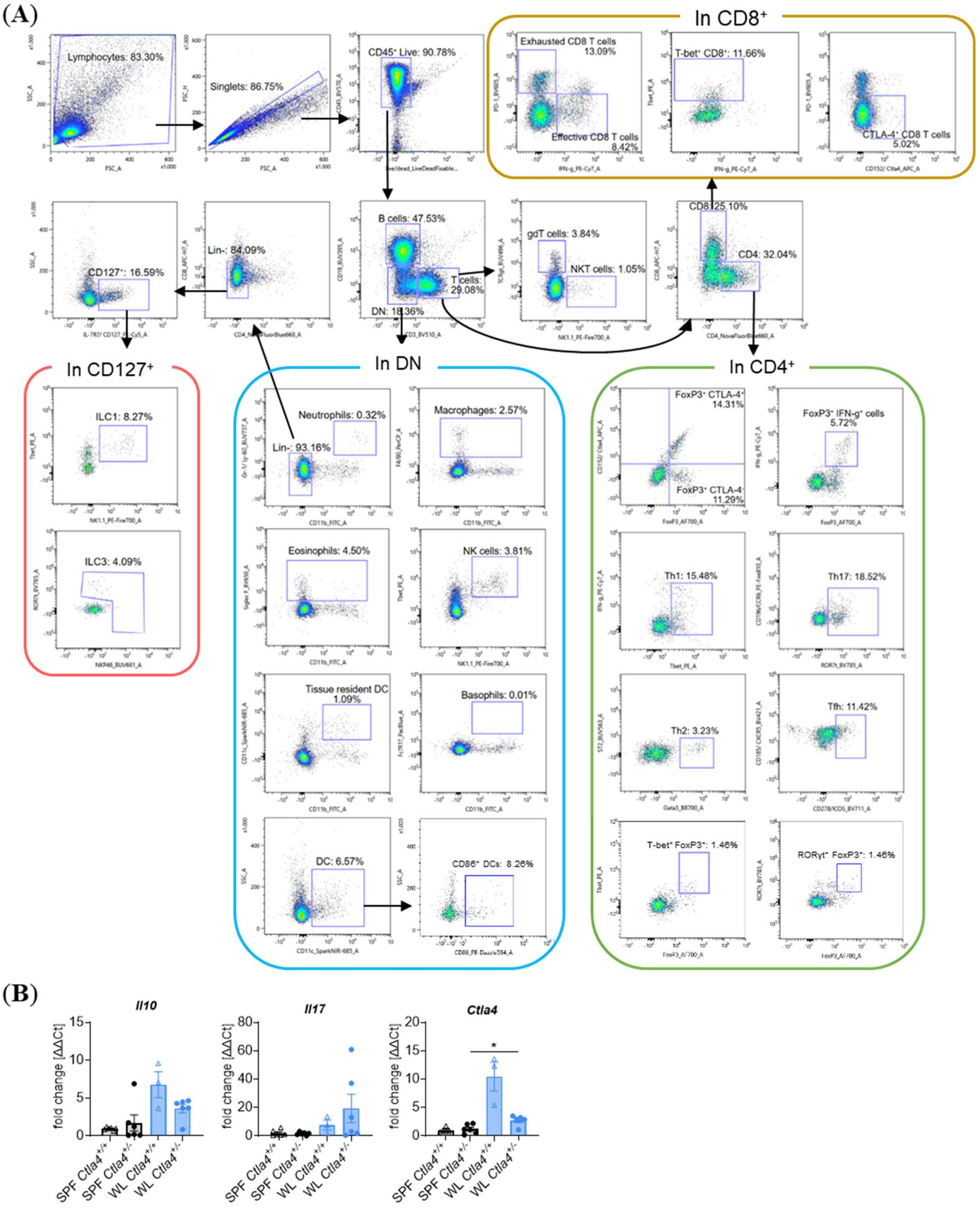


**Figure S5. Panel design and gating strategy for immune profiling.**

A. Flow cytometry gating scheme used to identify major immune cell populations analyzed in the study.

B. qPCR quantification of *Il10*, *Il17*, and *Ctla4* mRNA expression in colon tissue from 24-week-old SPF and WL mice, n=3-6. Data represent two independent experiments and were analyzed by unpaired t test. *p < 0.05, **p < 0.01.

(A)

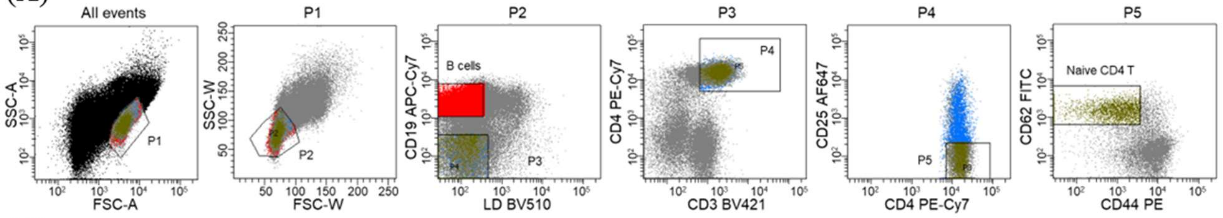
All events
P1
P2
P3
P4
P5
SSC-A
FSC-A
SSC-W
FSC-W
CD19 APC-Cy7
LD BV510
B cells
CD4 PE-Cy7
CD3 BV421
CD25 AF647
CD4 PE-Cy7
CD62 FITC
CD44 PE
Naive CD4 T

(B)

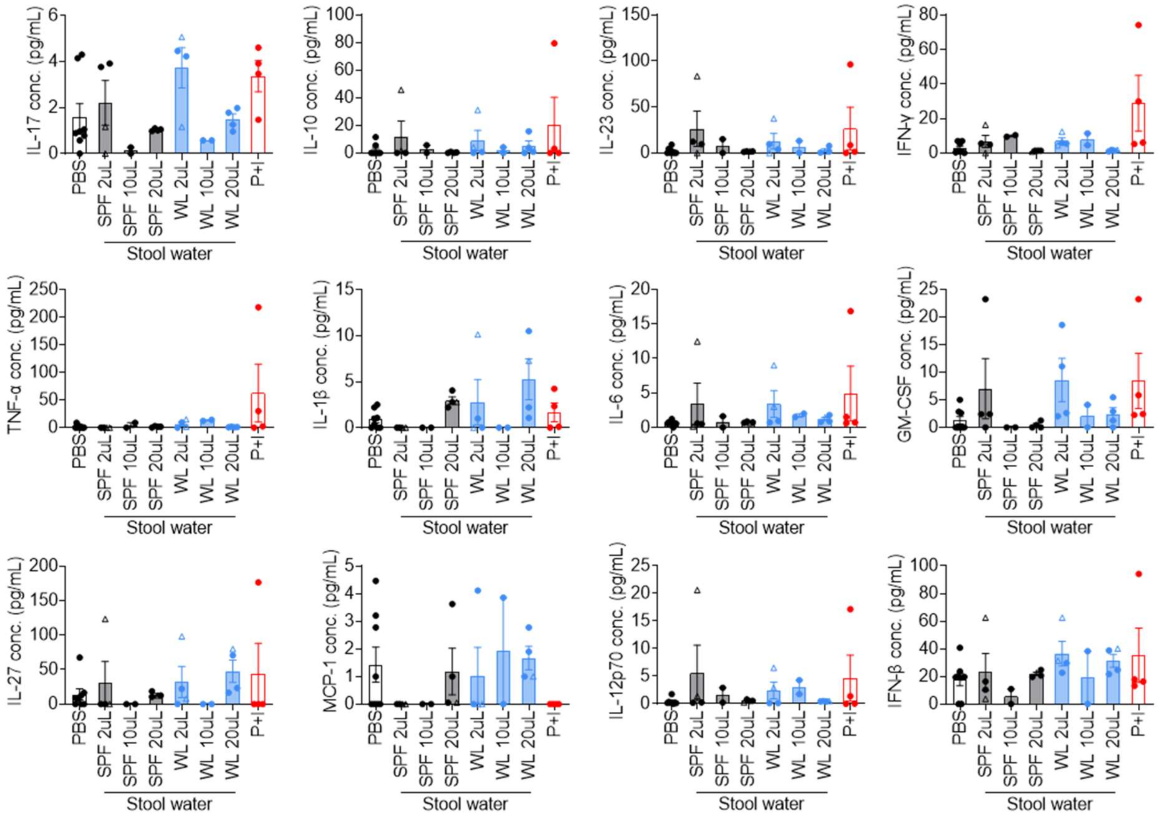
IL-17 conc. (pg/mL)
IL-10 conc. (pg/mL)
IL-23 conc. (pg/mL)
IFN-γ conc. (pg/mL)
TNF-α conc. (pg/mL)
IL-1β conc. (pg/mL)
IL-6 conc. (pg/mL)
GM-CSF conc. (pg/mL)
IL-27 conc. (pg/mL)
MCP-1 conc. (pg/mL)
IL-12p70 conc. (pg/mL)
IFN-β conc. (pg/mL)
PBS
SPF 2uL
SPF 10uL
SPF 20uL
WL 2uL
WL 10uL
WL 20uL
P+I
Stool water

(C)

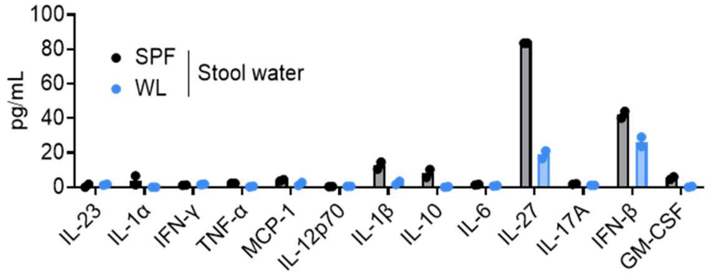
pg/mL
SPF
WL
Stool water
IL-23
IL-1α
IFN-γ
TNF-α
MCP-1
IL-12p70
IL-1β
IL-10
IL-6
IL-27
IL-17A
IFN-β
GM-CSF

(D)

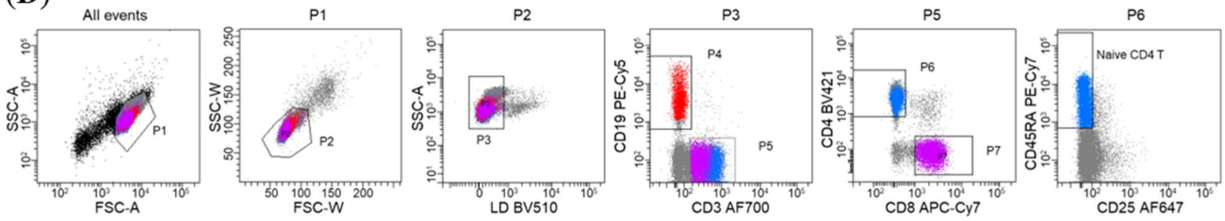
All events
P1
P2
P3
P5
P6
SSC-A
FSC-A
SSC-W
FSC-W
LD BV510
CD19 PE-Cy5
CD3 AF700
P4
P7
CD4 BV421
CD8 APC-Cy7
CD45RA PE-Cy7
CD25 AF647
Naive CD4 T

**Figure S6. Cytokine levels in stool water from SPF or WL mice.**

A. Gating strategy for sorting naïve CD4$^+$ T cells from mouse spleen.
B. Naïve splenic CD4$^+$ T cells derived from SPF *Ctla4*$^{+/-}$ mice were incubated for 16 hours with PBS, varying amounts of SPF or WL stool water from 11-week-old mice, and PMA+ionomycin (P+I). Culture supernatants were collected, and IL-1α concentrations were measured by ELISA. Circles and triangles represent responses induced by stool water from two different mice within each group, n=2-8.
C. Concentrations of multiple cytokines in stool water collected from 11-week-old SPF and WL *Ctla4*$^{+/-}$ mice were quantified using a LEGENDplex assay (n = 2).
D. Gating strategy for sorting naïve CD4$^+$ T cells from human PBMCs.

**Table S1. The scoring criteria for mouse disease severity.**

| | OBSERVATION | | POINT SCORING |
|---|---|---|---|
| COLITIS SPECIFIC SYMPTOMS | Weight reduction | 0-5% | 0 |
| | | >5-10% | 1 |
| | | >10-15% | 2 |
| | | >15-20% | 3 |
| | | >20-25% | 4 |
| | | > 25% | 10 (termination criterion) |
| | Stool consistency<br>Diarrhea | Normal | 0 |
| | | Soft stool | 2 |
| | | Watery diarrhea | 4 |
| | Blood in the stool | No blood | 0 |
| | | Slight bleeding | 2 |
| | | Severe bleeding | 10 (termination criterion) |
| NON-SPECIFIC SYMPTOMS | Fur | Normal | 0 |
| | | Coat defect (reduced grooming) | 2 |
| | Vocalization | None | 0 |
| | | Provoked | 2 |
| | | Unprovoked | 4 |
| | Eyelid slit | Open | 0 |
| | | Half open | 2 |
| | | Sunk and half closed | 4 |
| | Attitude | Normal posture | 0 |
| | | Abnormal posture | 1 |
| | Activity<br>Social behavior | Normal behavior/activity | 0 |
| | | Reduced activity / social contacts | 2 |
| | | Inactive / Self-isolation | 4 |
| | Piloerection | None | 0 |
| | | Piloerection | 1 |
| LOAD LEVEL | | MEASURES | TOTAL POINTS |
| Load level 0 = no load | | Monitoring three times a week | 0 |
| Load level 1 = low load | | Continue to monitor carefully; daily (on working days) visual monitoring of activity, posture and physical condition without touching or weighing the mouse. | 1-5 |

| | | |
|---|---|---|
| Stress level 2 = moderate stress | Continue to monitor carefully; daily checks including weighing and scoring, consult a vet if necessary and discuss further action. | 6-9 |
| Stress level 3 = high stress | Termination criterion: immediately terminate the experiment and euthanise the animal | ≥ 10 |

**Table S2. Mouse histology sample evaluation criteria.**

| Organ | Criteria | Score |
|---|---|---|
| **Liver** | Fibrosis | 0 – no fibrosis detectable; 1 - <20% in 10x overview of area; 2 – 21-50% in 10x overview of area; 3 - >50% in 10x overview of area |
| | Lymphocytic infiltration | 0 – no lymphocytic infiltration; 1 - <20% in 10x overview of area; 2 – 21-50% in 10x overview of area; 3 - >50% in 10x overview of area |
| | Lymph follicles | 0 – no follicle detectable at 10x; 1 – follicle detectable at 10x |
| | Steatosis | 0 – no steatosis detectable; 1 - <20% in 10x overview; 2 - >21% in 10x overview |
| **Small intestine & Colon** | Lymphocytic infiltration | 0 – no lymphocytic infiltration; 1 - <20% in 10x overview of area; 2 – 21-50% in 10x overview of area; 3 - >50% in 10x overview of area |
| | Granulocytes | 0 – no granulocytic infiltration; 1 - <20% in 10x overview of area; 2 – 21-50% in 10x overview of area; 3 - >50% in 10x overview of area |
| | Ulcerations | 0 – no ulcerations detectable at 10x; 1 – ulcerations detectable at 10x |
| | Erosions | 0 – no erosions detectable at 10x; 1 – erosions detectable at 10x |
| **Lung** | Lymphocytic infiltration | 0 – no lymphocytic infiltration; 1 - <20% in 10x overview of area; 2 – 21-50% in 10x overview of area; 3 - >50% in 10x overview of area |
| | Granulocytes | 0 – no granulocytic infiltration; 1 - <20% in 10x overview of area; 2 – 21-50% in 10x overview of area; 3 - >50% in 10x overview of area |
| | Fibrosis | 0 – no fibrosis detectable; 1 - <20% in 10x overview of area; 2 – 21-50% in 10x overview of area; 3 - >50% in 10x overview of area |
| | Hemorrhage | (0 – no hemorrhage detectable; 1 – hemorrhage detectable in 10x magnification |
| **Spleen** | Red pulp | 3 – normal distribution / presence in all fields of view; 2 – slight reduction up to 20% in fields of view; 1 – moderate reduction up to 50% in fields of view; 0 – strong reduction more than 50% in fields of view |
| | White pulp | 3 – normal distribution / presence in all fields of view; 2 – slight reduction up to 20% in fields of view; 1 – moderate reduction up to 50% in fields of view; 0 – strong reduction more than 50% in fields of view |
| | Hematopoiesis | 0 – no EMH detectable at 10x; 1 – EMH detectable at 10x |
| | Fibrosis | 0 – no fibrosis detectable; 1 - <20% in 10x overview of area; 2 – 21-50% in 10x overview of area; 3 - >50% in 10x overview of area |

| | | |
|---|---|---|
| **Skin** | Ulcerations | 0 – no ulcerations detectable at 10x; 1 – ulcerations detectable at 10x |
| | Erosions | 0 – no erosions detectable at 10x; 1 – erosions detectable at 10x |
| | IE Lymphocytes | 0 – no lymphocytic infiltration; 1 - <20% in 20x of area; 2 – 21-50% in 20x overview of area; 3 - >50% in 20x overview of area |
| | Fibrosis | 0 – no fibrosis detectable; 1 - <20% in 10x overview of area; 2 – 21-50% in 10x overview of area; 3 - >50% in 10x overview of area |
